# Pericyte-specific vascular expression of SARS-CoV-2 receptor ACE2 – implications for microvascular inflammation and hypercoagulopathy in COVID-19

**DOI:** 10.1101/2020.05.11.088500

**Authors:** Liqun He, Maarja Andaloussi Mäe, Lars Muhl, Ying Sun, Riikka Pietilä, Khayrun Nahar, Elisa Vázquez Liébanas, Malin Jonsson Fagerlund, Anders Oldner, Jianping Liu, Guillem Genové, Lei Zhang, Yuan Xie, Stefanos Leptidis, Giuseppe Mocci, Simon Stritt, Ahmed Osman, Andrey Anisimov, Karthik Amudhala Hemanthakumar, Markus Räsänen, Olivier Mirabeau, Emil Hansson, Johan Björkegren, Michael Vanlandewijck, Klas Blomgren, Taija Mäkinen, Xiao-Rong Peng, Thomas D. Arnold, Kari Alitalo, Lars I Eriksson, Urban Lendahl, Christer Betsholtz

**Author notes:** Shared contribution (Liqun He, Maarja Andaloussi Mäe, Lars Muhl, Ying Sun).

## Abstract

Accumulating clinical observations implicate vascular inflammation as an underlying cause of coagulopathy in severely ill COVID-19 patients and it was recently suggested that SARS-CoV-2 virus particles infect endothelial cells. Here, we show that endothelial cells do not express angiotensin-converting enzyme-2 (ACE2), the SARS-CoV-2 receptor. Instead, pericytes and microvascular smooth muscle cells express ACE2 in an organotypic manner. Pericyte deficiency leads to increased endothelial expression and release of Von Willebrand factor and intravascular platelet and fibrin aggregation, suggesting that pericytes limit endothelial pro-thrombotic responses. That pericytes and not endothelial cells express ACE2 may provide important clues to the pathology of COVID-19, as pericytes are normally shielded behind an endothelial barrier and may get infected only when this barrier is compromised by COVID-19 risk factors.

---

COVID-19 has currently been diagnosed in more than 11.0 million people worldwide and resulted in more than 530,000 deaths (as of July 5, 2020; Johns Hopkins University COVID-19 case tracker https://coronavirus.jhu.edu). COVID-19 is caused by the severe acute respiratory syndrome coronavirus-2 (SARS-CoV-2). Like other coronaviruses^1^, SARS-CoV-2 infects cells through binding of its spike (S) protein to cell surface receptors, followed by S protein cleavage, fusion of viral and cellular membranes and viral RNA entry into the cytoplasm. ACE2 has been identified as the cell surface receptor for SARS-CoV-2, and the proteases TMPRSS2 and cathepsin B/L (CTSB/L) have been shown to mediate SARS-CoV-2 S cleavage^2^. SARS-CoV-2 thereby appears to utilize the same molecular pathway for cellular entry as SARS–CoV-1, which caused the SARS epidemic in 2002-2003^3,4^. ACE2 normally functions in the renin/angiotensin pathway by cleaving angiotensin-2 to angiotensin 1-7, a peptide with multiple reported functions^5,6^.

Several types of epithelial cells in the nasal cavities, lung, gastrointestinal tract and eye were recently shown to express ACE2 and TMPRSS2 or CTSL, indicating susceptibility for SARS-CoV-2 infection^7–9^. These cellular targets offer explanation for the initial range of COVID-19 symptoms, including cough, fever, respiratory impairment, lost sense of smell and taste, nausea and diarrhea. Later in the disease course, however, some COVID-19 patients, experience a second round of severe symptoms and complications, including venous and arterial thrombosis with pulmonary embolism, myocardial infarction and stroke, acute kidney injury and neurological manifestations^10–16^. Moreover, some patients display severe hypoxemia without concomitant respiratory dysfunction early in the disease course^17^. These severe and partly paradoxical manifestations pose questions about possible vascular involvement secondary to the initial viral pneumonia. Laboratory blood tests reveal coagulation abnormalities that differ from those observed in certain other severe diseases. The COVID-19-associated coagulopathy (CAC) appears distinct from disseminated intravascular coagulation (DIC), for example^18–20^. In addition, severely ill COVID-19 patients display indicators of systemic inflammation^21,22^. The pathophysiological basis of these problems is not known, but involvement of the vascular endothelium has been suspected, posing the question if SARS-CoV-2 virus infects endothelial cells. Endothelial ACE2 expression has been reported based on immunodetection^5,23,24^, and recently the presence of SARS-CoV-2 virus particles in endothelial cells in COVID-19 patients was proposed^25,26^, but also questioned^27^. Recent single-cell RNA-sequencing (scRNAseq) studies have implicated low-level *ACE2* expression by endothelial cells across multiple organs^8^ but also vascular mural cells (pericytes and vascular smooth muscle cells (VSMC)) have been suggested to express ACE2 based on immunodetection^23^ and RNA sequencing^28–32^. However, these interpretations are clouded by ambiguities concerning antibody specificity and cell contaminations in scRNAseq data.

To know what (if any) vascular cell types express ACE2 is important for the understanding of COVID-19 vascular pathophysiology. If endothelial cells carry ACE2 receptors, their infection would likely only require viral dissemination from the primary infected cells into the blood (viremia), which has been reported for SARS-CoV-2^33,34^. However, infection of pericytes or other perivascular cells would require virus passage across the endothelial barrier, something that seems unlikely in a healthy vasculature given that corona virus particles are larger than the physiological endothelial pore size of most, if not all, blood vessels. However, in a number of pathophysiological conditions, including hypertension, inflammation, diabetes and obesity, the endothelial barriers are compromised and vascular permeability increased^35–39^. Notably, these conditions are also recognized as risk factors for severe COVID-19 disease^40,41^.

Here, we report that there is strong and specific ACE2 expression in microvascular mural cells in a highly organotypic fashion, while vascular endothelial cells are consistently ACE2 mRNA and protein negative. We also find that pericyte deficiency promotes endothelial synthesis and release of Von Willebrand factor, platelet aggregation and fibrin deposition, showing that pericyte injury may trigger endothelial pro-coagulant responses.

## Results

### *Ace2/ACE2* is specifically expressed in microvascular mural cells of the central nervous system

It is an emerging view that COVID-19 patients in hospital care often display CAC and systemic inflammation, both associated with poor prognosis^10,11,42–48^. To assess the pathology spectrum in a Swedish patient cohort, we studied 20 critically ill patients admitted to the intensive care unit at the Karolinska University Hospital, Stockholm, Sweden with positive diagnostic test for COVID-19. These patients uniformly displayed indicators of systemic inflammation, including markedly elevated C-reactive protein (CRP) and high circulating levels of pro-inflammatory cytokines, such as IL-6. They also displayed a typical CAC biochemistry and symptomatology, including elevated D-dimer levels and pulmonary embolism despite prophylactic anticoagulation therapy (Extended Data Table 1).

To explore the molecular basis for a direct infection of vascular cells by SARS-CoV-2 as a putative underlying case of CAC and systemic inflammation, we analyzed the expression of ACE2 and S priming proteases by vascular cells. Mice and humans frequently share gene and protein expression patterns across cell types, and we therefore first assessed the expression and distribution of *Ace2/ACE2* in the mouse vasculature. Using our previously published scRNAseq database of adult mouse brain vascular gene expression patterns^29,30^, we found that *Ace2* mRNA is highly enriched in pericytes and venous VSMC, qualifying *Ace2* among the top 15 specific markers for these cells (Extended Data Fig 1a,b and http://betsholtzlab.org/VascularSingleCells/database.html). At lower levels, *Ace2* was also found in VSMCs negative for *Cnn1* and other arterial VSMC markers (Extended Data Fig 1c,d). A subpopulation of *Ace2*-positive type-2 fibroblasts (FB2) cells and a few *Ace2*-positive endothelial cells were noted, however these cells coexpressed *Kcnj8* (Extended Data Fig 1c) and several other pericyte markers (Extended Data Fig 1c, e), suggesting that they were pericyte-contaminated. Deep analysis of the 23 *Ace2*-positive endothelial cells in this dataset revealed that 20 expressed at least one additional pericyte marker and 18 expressed multiple pericyte markers, whereas these markers were generally absent from *Ace2*-negative endothelial cells (Extended Data Fig 1e). We failed to find any distinguishing gene expression pattern beyond the addition of pericytes markers onto the transcriptomes of FB2 and endothelial cells, respectively (Extended Data Fig 1f-h), together arguing that the presence of *Ace2* RNA sequences in FB2 and endothelial cells is caused by pericyte contamination.

To deepen the analysis of brain vascular cells, we devised a computational pipeline for meta-analysis of scRNAseq data generated using SmartSeq2 and DropSeq (10x Genomics) platforms (see Methods). We used this pipeline to merge two published^29,30,49^ and one unpublished dataset generated using both platforms. The combined data are available for analysis gene-by-gene at http://betsholtzlab.org/Publications/BrainIntegration/search.html (login ID: reviewer, password: reviewer). This analysis confirmed that *Ace2* is specifically expressed in brain pericytes, venous VSMCs and *Cnn1*-negative arteriolar VSMCs (Fig 1a-c). An additional *Ace2*-positive cell cluster contributed uniquely from the DropSeq dataset (cluster #17 in Fig 1a-c) contained both pericyte and endothelial markers but lacked other unique transcripts, and we therefore concluded that it was composed of pericyte-contaminated endothelial cells.

**Figure 1:**
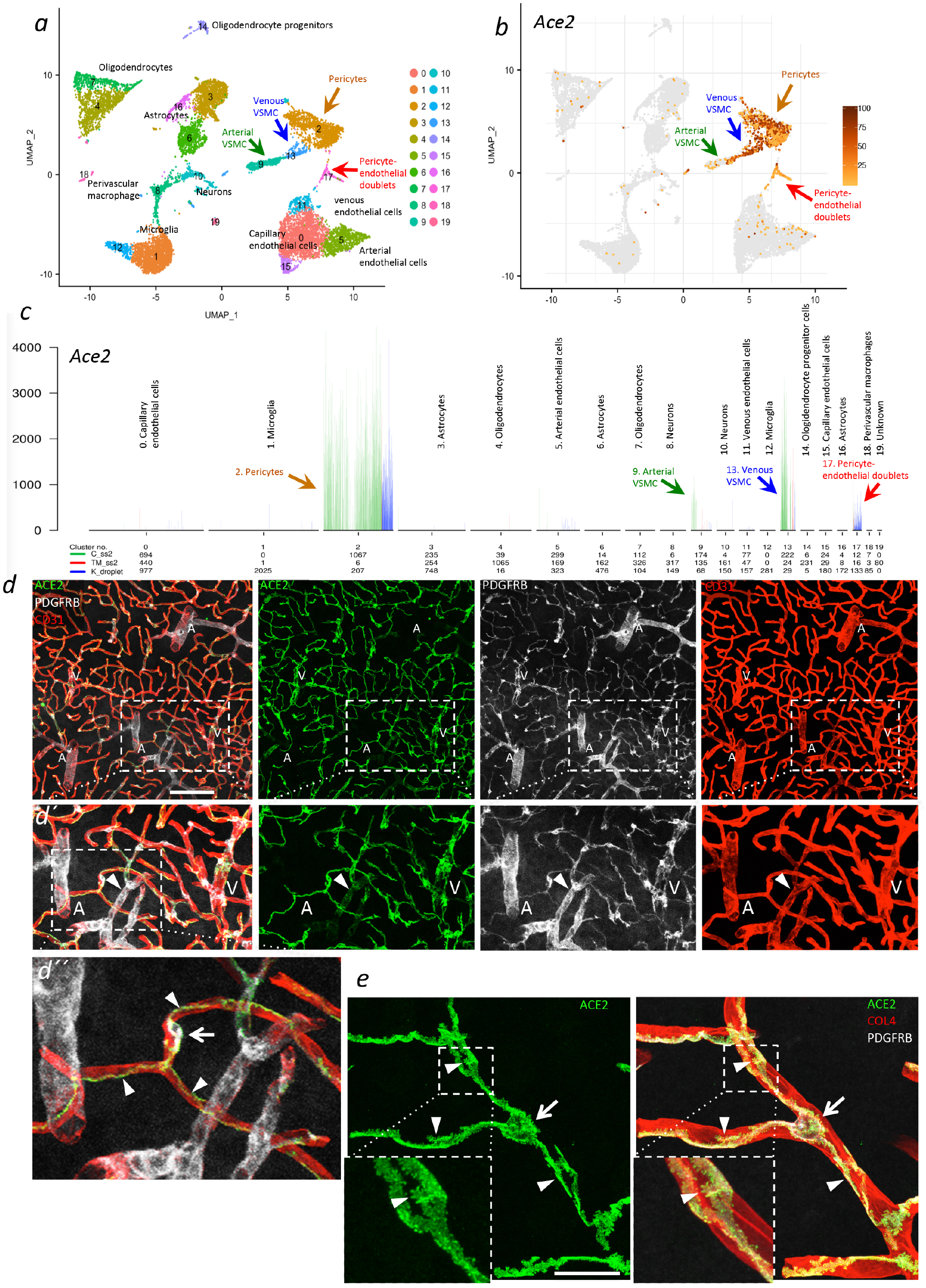
ACE2 expression in adult mouse brain. ***a*,** UMAP display of integrated mouse brain scRNASeq data. Coloring is based on cluster assignment and cellular annotations are based on canonical marker expression available at http://betsholtzlab.org/Publications/BrainIntegration/search.html ***b*.** The same UMAP display as in ***a*** with *Ace2* expression overlay (dark color represents higher expression and grey color represents *Ace2*-negative cells). ***c*.** Bar plots of the normalized expression levels of *Ace2* in each cluster. Cell type annotations for each cluster are indicated. Individual bars represent single cells and are colored according to the data source (see Methods) as indicated together with cell numbers contributions below the x-axis. Arrows of different colors in ***a-c*** indicate the corresponding *Ace2*-positive clusters in the UMAP and bar plot displays. ***d,e*.** IF detection of ACE2 in adult mouse brain in combination with the indicated markers. Insets indicate magnified areas. Left-right shows the same microscopic field with different antibody stainings. CD31 and COL4 are used as markers for the endothelium and basement membrane, respectively. PDGFRB marks mural cells. Note the similar expression of ACE2 and PDGFRB in pericytes and venous/venular VSMC, the absence of ACE2 from large arterioles and weak presence at terminal arterioles (arrowhead in ***d’***). In ***d”*** and ***e***, the arrows point at the pericyte cell soma and the arrowheads at its primary processes. In the insets in e, the arrowhead point at the pericyte’s secondary processes. A, arteriole; V, venule. Scale bars: 100 μm in ***d*** and 10 μm in ***e*.**

We next assessed the localization of ACE2 protein-expressing cells in the mouse brain vasculature by immune-fluorescence (IF) analysis. In agreement with the scRNAseq data, we found the brain ACE2 IF signal was concentrated to peri-endothelial cells with typical morphology of pericytes: a distinct cell soma, primary longitudinally oriented processes and secondary short club-like extensions (Fig 1 d,e). The endothelial cells were invariably ACE2 IF negative. As expected for a cell membrane-bound protein, ACE2 showed a similar sub-cellular distribution as platelet-derived growth factor receptor-beta (PDGFRB) and N-aminopeptidase (CD13), (Fig 1d and Extended Data Fig 2a) and a slightly distinct localization compared to the cytoplasmic *Pdgfrb-GFP* reporter (Extended Data Fig 2b). Focusing on the cortex where the arterio-venous vascular hierarchy is readily distinguishable by morphology and marker expression^29^, we found strongly ACE2-positive mural cells associated with capillaries and veins/venules (Fig 1d) and weakly ACE2-positive VSMCs at terminal arterioles (Fig 1d and Extended Data Fig 2c), whereas calponin (CNN1)-positive mural cells around larger arteries/arterioles were ACE2-negative (Fig 1d and Extended Data Fig 2c). This ACE2 IF pattern thus matched the zonation of *Ace2*-positive brain mural cells predicted from the scRNAseq data, revealing ACE2 as more specific to micro-vascular mural cells than other commonly used protein markers for mural cell (e.g. PDGFRB, CD13, and NG2), which also label arterial VSMC^50^.

To investigate the generality of the microvascular pattern of mural cell ACE2 expression in the central nervous system (CNS), we additionally analyzed spinal cord and eye. Similar to the brain, spinal cord ACE2 was concentrated to cells with the typical morphology and marker expression of pericytes (Extended Data Fig 2d) and weaker ACE2 expression was noted also in alpha-smooth muscle actin (αSMA)-positive VSMCs at terminal arterioles, whereas CNN1-positive VSMCs of larger arteries were ACE2-negative (Extended Data Fig 2e,f). Analysis of the eye (Extended Data Fig 3a-t) showed that also retinal pericytes were strongly ACE2-positive (Extended Data Fig 3c,g,k,o) and αSMA–positive VSMC in small diameter retinal arterioles were weakly ACE2-positive, whereas larger diameter arterioles/arteries were ACE2-negative (Extended Data Fig 3k). Also the choriocapillaris harbored strongly ACE2-positive pericytes (Extended Data Fig 3q-t), whereas αSMA–positive vessels feeding this capillary plexus were ACE2-negative (Extended Data Fig 3c,k,s). ACE2-positive pericytes were further found in the ciliary body (Extended Data Fig 3h). Together, these analyses reveal a common pattern and zonation of ACE2 expression in CNS mural cells. Endothelial cells were consistently ACE2-negative at all analyzed locations. Intriguingly, pericytes located in the extra-ocular skeletal muscle, hence residing outside of the CNS, were ACE2-negative (Extended Data Fig 3d), demonstrating that mural cell ACE2 expression is highly organotypic. We further noted that the surface epithelium of the conjunctiva and cornea was ACE2-positive (Extended Data Fig 3g), confirming recent observations by others^8^.

To extend the analysis of *Ace2* expression to non-vascular cell types in several regions of the brain, we explored the mousebrain.org atlas, which confirmed that *Ace2* is not appreciably expressed outside of the vasculature across >250 different central and peripheral nervous system cell types^51^ (http://mousebrain.org/genesearch.html). More specifically, the mousebrain.org atlas report *Ace2* mRNA in cells annotated as pericytes, but also in a cluster of endothelial cells^51^, which, however, we found to co-express numerous pericyte markers (including *Pdgfrb* and *Notch3*) suggesting contamination. The mousebrain.org atlas denotes weak *Ace2* expression also in neurons in dorsal root ganglia^51^.

To analyze *ACE2* expression in mural cells of the human CNS is currently challenging, given the scarcity of scRNAseq data for these cells. However, in a dataset representing the developing human prefrontal cortex^52^, we identified six *ACE2*-positive cells expressing at least one pericyte marker, and three of them expressed multiple pericyte markers (Extended Data Fig 4a). We also assessed a human glioblastoma Drop-Seq (10 x Genomics) scRNAseq dataset consisting of 69,125 endothelial and immune cells sorted using anti-CD31 antibodies (Zhang et al, unpublished). Among these, we found only two *ACE2*-positive cells, both of which had pericyte markers (Extended Data Fig 4b), suggesting that the *ACE2* mRNA sequences were derived by pericyte contamination, and that endothelial cells were *ACE2*-negative. Finally, we assessed the scRNAseq data from an analysis of human retinas^53^ and found enrichment for *ACE2* in pericytes as compared to other retinal cells types, including endothelial cells (data not shown). Together, these observations indicate that pericytes express *ACE2* in the human brain, but additional analysis are needed to confirm these conclusions. We failed to find any indications for endothelial ACE2 expression in the analyzed human datasets.

### *ACE2* is specifically expressed in pericytes of the heart

We next analyzed *Ace2/ACE2* expression in mouse and human heart. Based on scRNAseq and single-nucleus (sn)RNAseq data, it was recently suggested that *ACE2* expression occurs across multiple human cardiac cell types, including cardiomyocytes, endothelial cells, pericytes, fibroblasts and macrophages^8,31,32^. When examining adult mouse heart scRNAseq data enriched for mesenchymal cells collected by fluorescence-activated cell sorting (FACS) from *Pdgfrb-GFP* transgenic mice (Muhl et al, in press), we found prominent expression of *Ace2* in pericytes, whereas we could not detect *Ace2* mRNA in fibroblasts and VSMC (data not shown). To expand this analysis to additional cardiac cell types, we used our meta-analysis pipeline to integrate three unpublished and one published^49^ datasets. These combined data, available for gene-by-gene search at http://betsholtzlab.org/Publications/HeartIntegration/search.html, (login ID: reviewer, password: reviewer) provided comprehensive coverage of the principal cell types in the mouse heart, including cardiomyocytes, different types of endothelial cells (blood vascular and endocardial), mural cells (pericytes, VSMCs), and subtypes of fibroblasts and macrophages (Fig 2a). Of these, only pericytes expressed *Ace2* substantially (Fig 2b, c). As in the brain, rare endothelial cells displayed RNAseq counts for *Ace2*, however, out of 21 cells in total, 18 expressed at least one pericyte marker and 15 expressed multiple pericyte markers (Extended Data Fig 5a,b), in contrast to *Ace2*-negative endothelial cells (Extended Data Fig 5b,c), together suggesting that the rare *Ace2*-sequences found in cardiac endothelial cells were contributed by pericyte-contamination.

**Figure 2:**
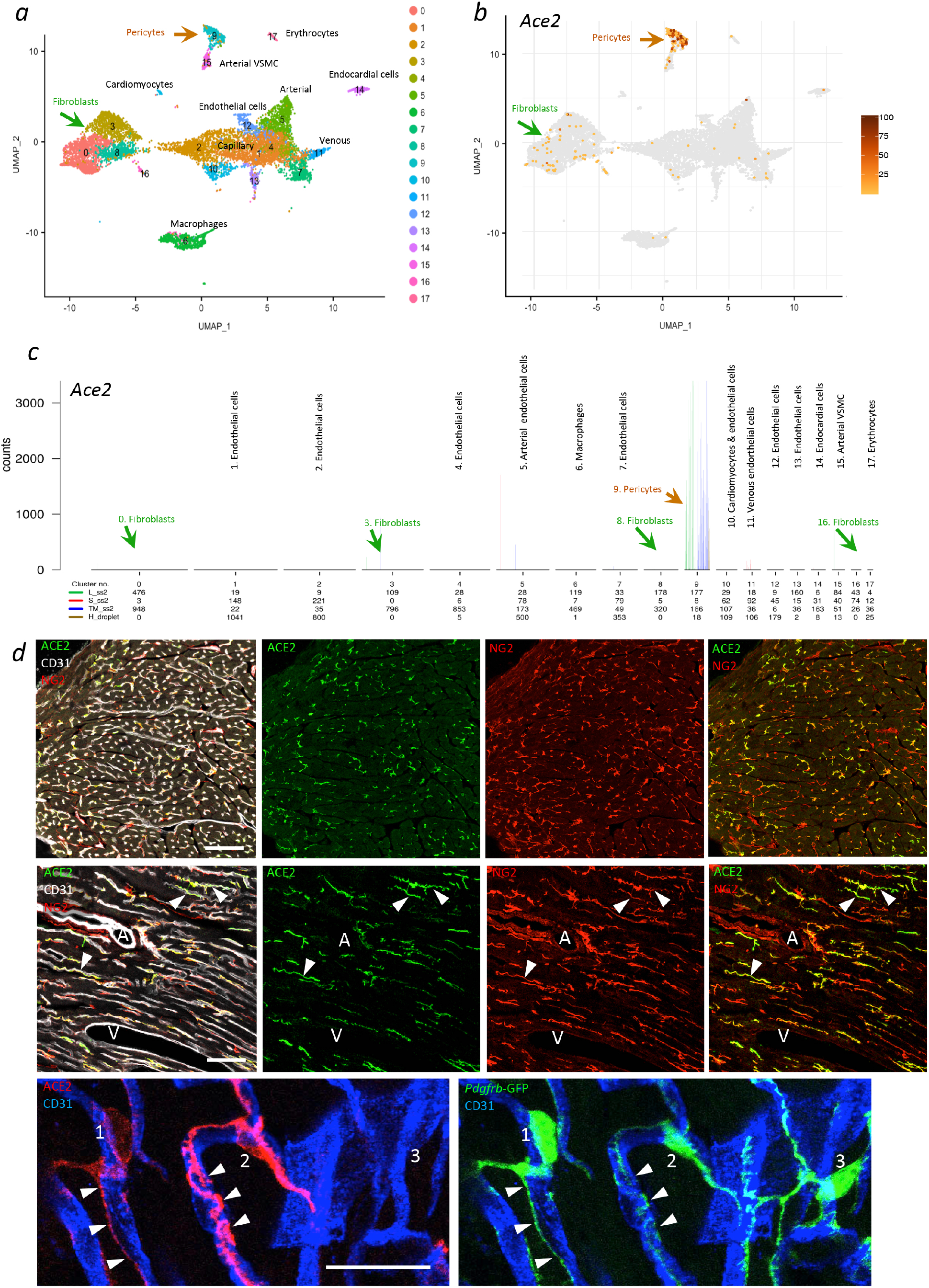
ACE2 expression in the adult mouse heart. ***a*.** UMAP display of integrated mouse heart scRNASeq data. Coloring is based on cluster assignment and cellular annotations are based on canonical marker expression available at http://betsholtzlab.org/Publications/HeartIntegration/search.html. ***b*.** The same UMAP display as in ***a*** with *Ace2* expression overlay (dark color represents higher expression and grey color represents *Ace2*-negative cells). ***c*.** Bar plots of the normalized expression levels of *Ace2* in each cluster. Cell type annotations for each cluster are indicated. Individual bars represent single cells and are colored according to the data source (see Methods) as indicated together with cell numbers contributions below the x-axis. Arrows of different colors in ***a-c*** indicate the corresponding *Ace2*-positive clusters in the UMAP and bar plot displays. ***d*.** IF detection of ACE2 in adult mouse brain in combination with the indicated markers. Each row of images shows the same microscopic field with different combinations of label, as indicated. CD31 is used as a marker for endothelium. NG2 and *Pdgfrb*-GFP marks mural cells. Note the overlapping expression of ACE2 and NG2 in a proportion but not all pericytes and that arterial and venous VSMC are ACE2-negative. The bottom row shows an example of three neighboring pericytes with different levels of ACE2 expression. Note also the different subcellular distribution of cell membrane associated ACE2 and cytoplasmic GFP expressed from the *Pdgfrb* promoter. A, artery; V, vein Scale bars: 100 μm, 50 μm and 25 μm respectively in top, middle and lower rows of images.

Cardiac ACE2 protein IF signal was detected only in cells with the typical location and morphology of pericytes: a round cell body and long processes adherent to the endothelial cells (Fig 2d). The ACE2 IF signal overlapped with *Pdgfrb*-GFP, albeit with the expected difference in subcellular localization: ACE2 in cell membrane and processes and GFP in cytoplasm and nucleus. While ACE2 IF signal was thus confined to pericytes in both CNS and heart, we noticed two differences between these organs. First, in heart *Ace2* mRNA and protein was found only in capillary pericytes (Fig 2c, d), whereas in CNS small arteriolar and venous mural cells were also positive. Second, not all cardiac capillary pericytes were strongly ACE2 positive; some were weakly positive and some were negative (Fig 2d).

In order to assess *ACE2* expression in the human heart, we re-analyzed scRNAseq data from healthy adult human hearts^54^ identifying a cluster of cells with substantially higher *ACE2* expression compared to other clusters (Extended Data Fig 6a,b). Based on canonical markers and inference from gene expression patterns in mouse where the transcriptomes of pericytes, VSMCs and fibroblasts have been deeply analyzed (Muhl et al, in press), the *ACE2*-high cluster was unambiguously identified as cardiac pericytes (Extended Data Fig 6b). A second cluster identified as fibroblasts contained a minor proportion of *ACE2*-positive cells (cluster 5 in Extended Data Fig 6b). One of the endothelial clusters contained a few *ACE2*-positive cells (cluster 2 in Extended Data Fig 6b), which, however, also expressed pericyte markers and were therefore concluded to be pericyte-contaminated (Extended Data Fig 6c). All other cardiac cells were *ACE2*-negative with the possible exception of cardiomyocytes, which although displaying lower *ACE2* levels than pericytes did not show signs of pericyte contamination. Collectively, our analysis of scRNAseq data from mouse and human healthy hearts establishes pericytes as the major cellular source of *Ace2/ACE2* in the adult heart, with putative low expression also in human fibroblasts and cardiomyocytes. ACE2 expression was consistently undetectable in cardiac endothelial cells in both mouse and human.

### In the lung, airway epithelial cells form the major *Ace2/ACE2* expression site

Lung epithelial cells are known primary targets for SARS-CoV-2 infection, but it has recently been proposed that also lung endothelial cells may become virus infected^26,55^. Our previously published scRNAseq dataset of pulmonary vascular cells^29^ did not reveal *Ace2* expression in lung vascular cells but showed a distinct *Ace2* signal in epithelial cells displaying markers of multiciliation (http://betsholtzlab.org/VascularSingleCells/database.html}. To provide deeper insight into the *Ace2* expression by different pulmonary cell types, we used our meta-analysis pipeline to integrate three published^29,30,49^ and two unpublished adult mouse lung scRNAseq datasets (Fig 3a). This analysis is available for gene-by-gene browsing at http://betsholtzlab.org/Publications/LungIntegration/search.html (login ID: reviewer, password: reviewer). The *Ace2* mRNA distribution in this combined data showed significant expression only in alveolar type-2 (AT-2) (surfactant protein C (*Sftpc*)-*positive*) and multiciliated airway epithelial cells (*Foxj1*-positive) (Fig 3a-c). No significant *Ace2* expression was observed in any of the subtypes of vascular endothelial cells (*Pecam1* positive), mural cells (*Notch3* positive), fibroblasts (*Pdgfrb* positive, *Notch3* negative), alveolar macrophages or other hematopoietic cells (*Ptprc* positive) (Fig 3c). A strong ACE2 IF signal was observed in the bronchial epithelium throughout the bronchial tree (Fig 3d). In the alveolar region distal to the terminal bronchioles, we found ACE2 IF signal in SFTPC-positive AT-2 cells (Fig 3d). We failed to find ACE2 in any endothelial populations in the lung, including alveolar capillaries and large vessels (Fig 3d and Extended Data Fig 7). Also CD68-positive alveolar macrophages were ACE2-negative (Extended Data Fig 7). While the vast majority of lung pericytes were ACE2 IF negative, particularly in the alveolar region, we found a few ACE2 positive pericytes close to larger bronchi and abundantly in trachea capillaries (Extended Data Fig 7).

**Figure 3:**
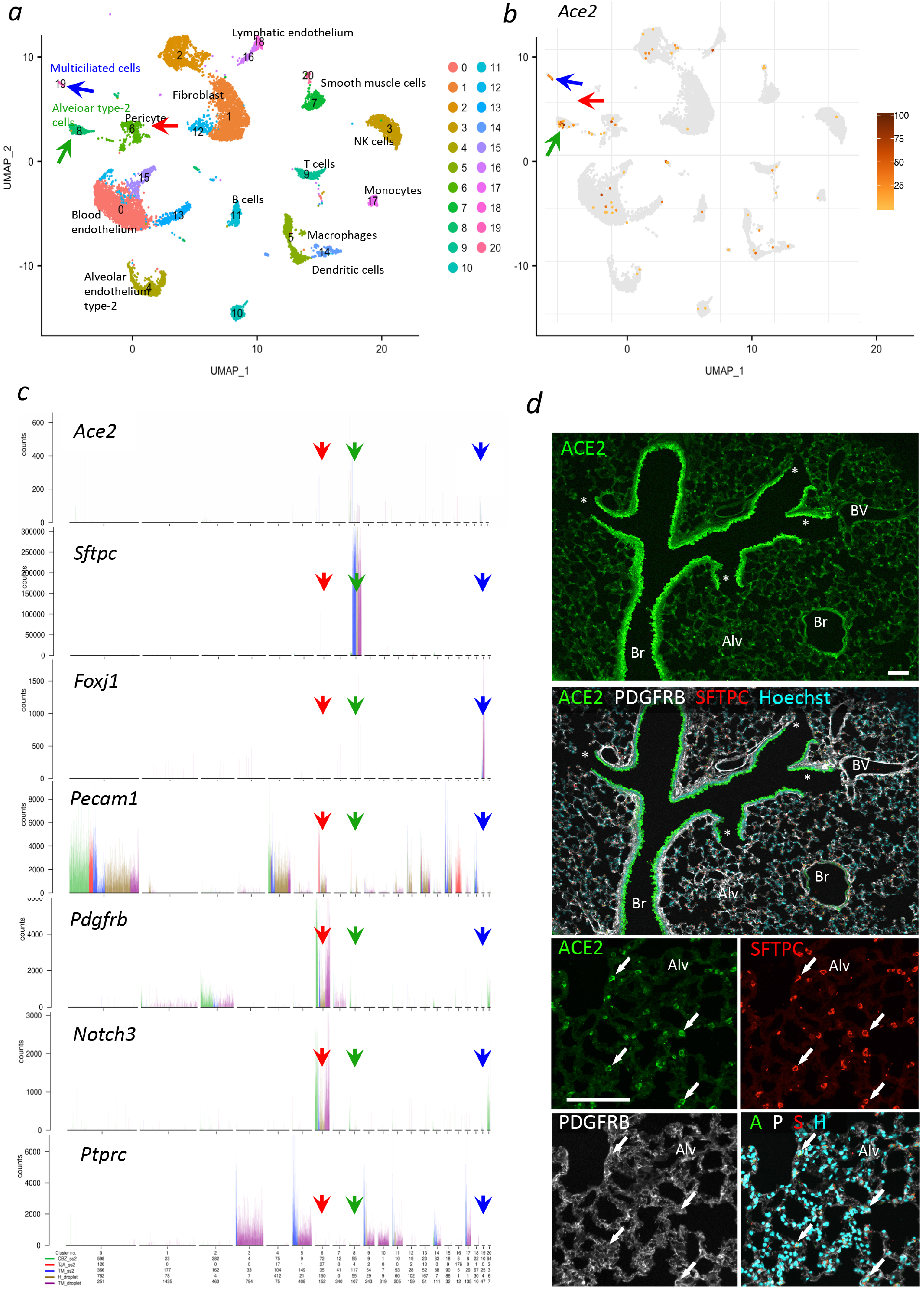
ACE2 expression in adult mouse lung. **a.** UMAP visualization of the integrated scRNASeq data from mouse lung (left) with *Ace2* expression overlay (right, darker color represents higher expression and grey color represents *Ace2*-negative cells). Colors indicate cluster assignment. Annotations were based on the expression of canonical markers for each indicated cell type available at http://betsholtzlab.org/Publications/LungIntegration/search.html. ***b***. The same UMAP display as in ***a*** with *Ace2* expression overlay (dark color represents higher expression and grey color represents *Ace2*-negative cells). For all panels, green arrows point at the AT-2 cell cluster, blue arrows on the multiciliated cell cluster and red arrows at the pericyte cluster. ***c***. Bar plots depict the expression of indicated genes across clusters. Each bar represents a single cell and is colored according to the indicated data source as indicated together with cell numbers below the bottom x-axis (see Methods). **c**. IF staining for indicated proteins in adult mouse lung. Prominent ACE2 signal is observed in bronchial epithelium. Asterisks mark end of terminal bronchioles. The lower panels show the same microscopic field of the alveolar region with different labels visualized. Arrows provide land marks and point at four AT-2 cells as indicated by the SFTPC staining. Note the overlap between ACE2 and SFTPC in the alveolar region and lack of ACE2 staining of pericytes (labeled by PDGFRB). No ACE2 IF signal was observed in endothelial cells. Br: bronchi, BV: blood vessel. Alv: alveolar region Scale bars: 100 μm.

In order to assess *ACE2* expression in human lungs, we re-analyzed scRNAseq data from the lungs of adult human transplant donors and lung fibrosis patients^56^. Here, AT-2 cells constituted the major cellular source of *ACE2* transcripts, but expression was detected also in basal, club and multiciliated cells (Extended Data Fig 8). Immune cells, endothelial cells and fibroblasts were *ACE2*-negative, but the vascular cell number was too small for a comprehensive analysis. Mural cells, for example, were not present in the dataset.

We also analyzed the expression of the SARS-CoV-2 S protein priming proteases in brain, heart and lung (Extended Data Fig 9). We observed *Ace2* and *Tmprss2* co-expression in epithelial cells across the mouse lung dataset, whereas *Ctsl* and *Ctsb* were more broadly expressed (albeit weakly in hematopoietic cells) (Extended Data Fig 9). However, *Tpmrss2* was not expressed in brain and heart pericytes, which instead exhibited co-expression of *Ace2* with *Ctsb* and *Ctsl* (Extended Data Fig 9).

Taken together, these observations reveal that in contrast to brain and heart and similar to ocular muscle, most vascular mural cells in the lung do not express ACE2. Instead our data show that in both mouse and human lung bronchial epithelial cells and alveolar AT-2 cells are the primary expression sites for ACE2 mRNA and protein. Furthermore, our data reveal differences in the S protein priming proteases that are expressed in pulmonary epithelial cells (TMPRSS2) versus brain and heart pericytes (CTSB/L).

### Pericyte hypoplasia elicits an endothelial pro-coagulant response

We identify pericytes as the predominant *Ace2*-expressing cells in CNS and heart. Given the roles of pericytes in vascular barrier formation and in restraining pro-inflammatory responses in endothelial cells^57–60^ (Mäe et al, submitted), we analyzed pro-thrombotic responses in adult mice with constitutive hypoplasia of pericytes caused by decreased PDGF-B signaling via PDGFR-β (*Pdgfb^ret/ret^* mice)^61^. In addition to the pericyte loss and development of dilated capillaries reported previously for the *Pdgfb^ret/ret^* brain^57^, we found increased levels of Von Willebrand Factor mRNA (*Vwf*) and protein (VWF) compared to *Pdgfb^ret/+^* littermate control mice, which had normal microvascular pericyte coverage (Fig 4a,b). Increased capillary VWF expression was observed also in *Pdgfb^ret/ret^* heart in comparison with *Pdgfb^ret/+^* littermate controls (Fig 5), which correlated with decreased pericyte coverage also in this organ (Extended Data Fig 10). VWF promotes platelet adhesion and blood coagulation in wounds through binding and stabilization of factor VIII. In controls, VWF mRNA and protein were mainly confined to arterioles and venules with limited or undetectable presence in capillaries, whereas the dilated capillaries showing signs of pro-inflammatory activation in *Pdgfb^ret/ret^* mice^57^ (Mäe et al, submitted) were strongly and uniformly VWF-positive (Fig 4a,b). The VWF IF signal was primarily localized to intracellular rod-shaped vesicles (Weibel-Palade bodies) (Extended Data Fig 10, Extended Data 11 and Extended Data Video 1), but it was also frequently observed as a “halo” around vessels in the *Pdgfb^ret/ret^* brain parenchyma, presumably due to local VWF release (Fig 4b,c). We next assessed platelet adherence and aggregation in the *Pdgfb^ret/ret^* vessels by analyzing CD41 IF and fibrinogen (FBG) leakage and fibrin deposition (Extended Data Fig 12). Sites displaying both platelet aggregation and fibrin deposition were commonly observed in brain sections from *Pdgfb^ret/ret^* mice (Extended Data Fig 12), whereas in controls, we observed only rarely individual adhered platelets, but no signs of platelet aggregation or FBG/fibrin deposition (data not shown). We also found that VWF release coincided with 70 kDa dextran tracer leakage into the brain parenchyma, demonstrating a local impairment of the blood-brain barrier at these sites (Extended Data Fig 12). In summary, these data reveal an important role for pericytes in restricting pro-coagulant responses concomitant with vascular leakage in endothelial cells.

**Figure 4.**
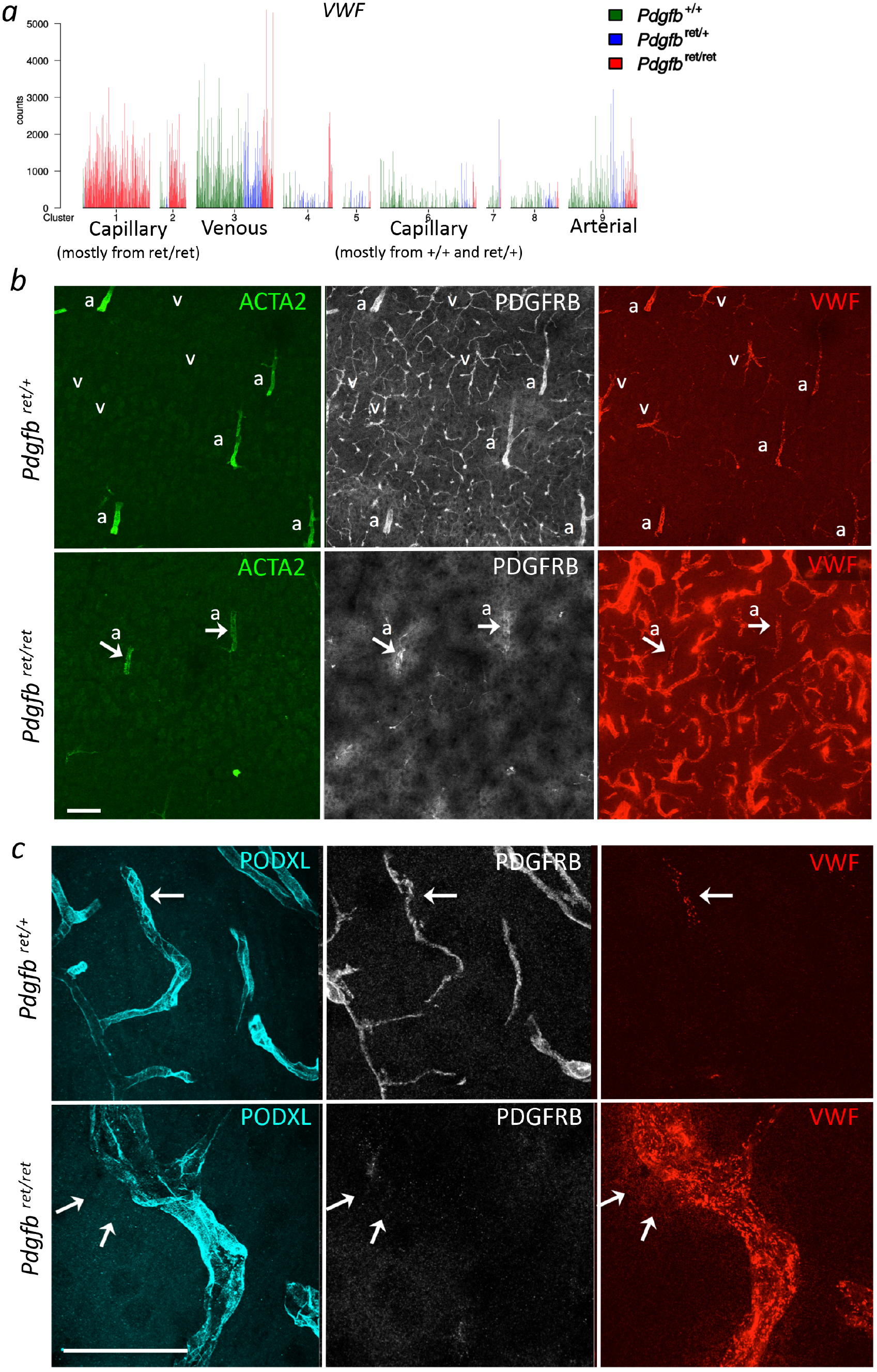
Increased Von Willebrand Factor expression in *Pdgfb^ret/ret^* brain. ***a*.** Bar plot illustrating *Vwf* expression in endothelial cells from *Pdgfb*^+/+^, *Pdgfb*^ret/+^ and *Pdgfb*^ret/ret^ brains. ***b-c*.** IF staining of mouse cortex vasculature in *Pdgfb*^ret/+^ and *Pdgfb*^ret/ret^ mice with antibodies against (***b***) VWF, PDGFRB and ACTA2, and (**C**) PODXL, PDGFRB and VWF). a, arterioles. v, venules. Note that in controls, capillaries covered by PDGFRB-positive pericytes are VWF-negative, whereas the dilated pericyte-deficient capillaries in *Pdgfb*^ret/ret^ brain are strongly VWF-positive. *c*. Normal brain capillaries (PODXL labels endothelial cells) covered by PDGFRB-positive pericytes are essentially VWF-negative (arrow points at location of a few Weibel-Palade bodies). In contrast, the pericyte-deficient capillaries in *Pdgfb*^ret/ret^ brain show dense content of Weibel-Palade bodies and local “halo” (arrows) of VWF released into the tissue interstitium. Scale bars: 50 μm.

**Figure 5.**
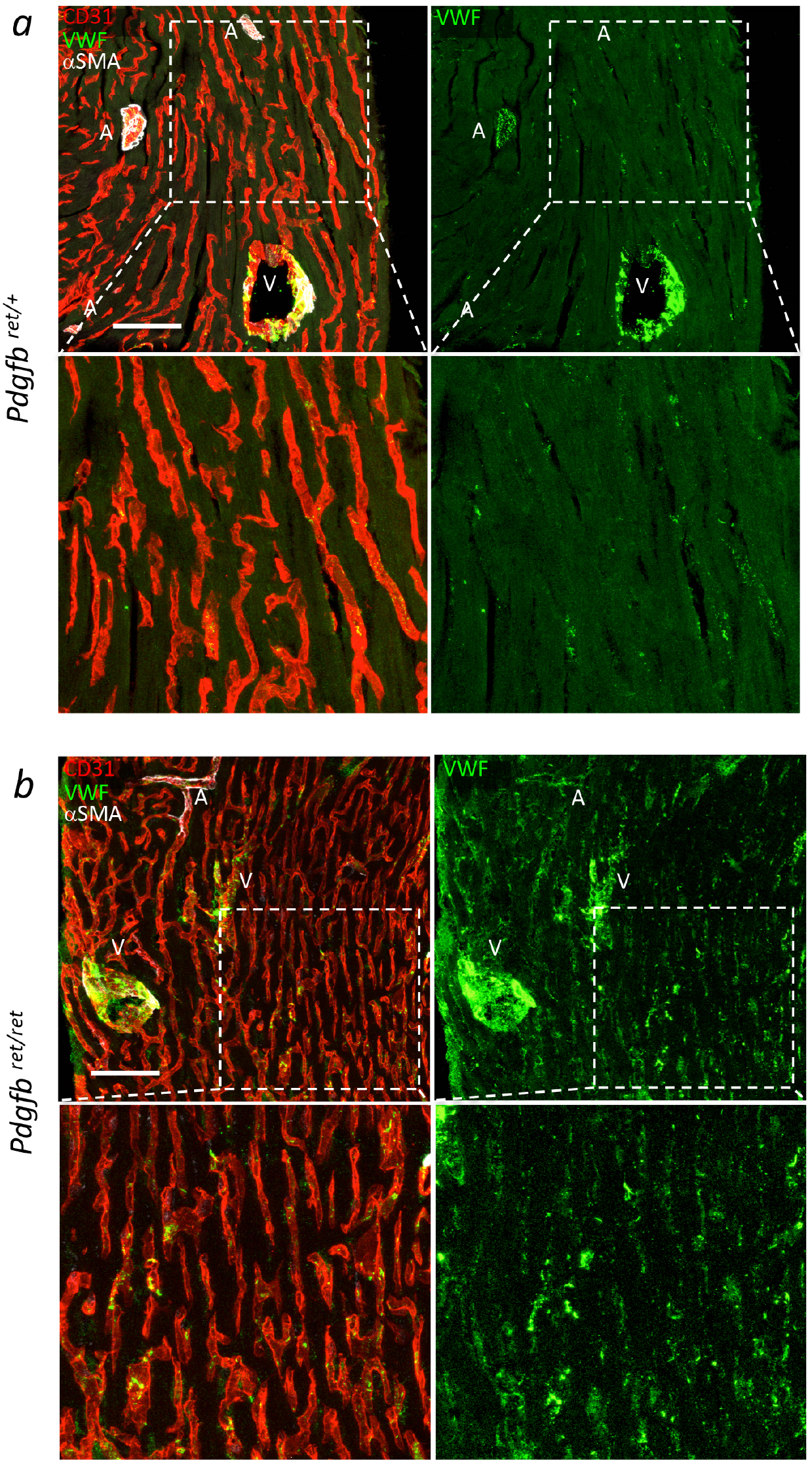
Increased Von Willebrand Factor expression in *Pdgfb^ret/ret^* heart. Confocal microscopy images of sections from adult control *Pdgfb*^ret/+^ (***a***) and mutant *Pdgfb*^ret/ret^ (***b***) mouse hearts IF stained with the indicated antibodies. Horizontal image pairs depict the same microscopic field with different label. Note the VWF expression in control veins and arteries, with few CD31 positive capillary endothelial cells being VWF positive. In pericyte-deficient *Pdgfb*^ret/ret^ hearts, VWF expression is increased in capillary endothelium (compare insets in the upper and lower panels). A: artery/arteriole, V: vein. Scale bars: 50 μm.

## Discussion

### The “COVID-19-pericyte hypothesis”

Our results lead us to posit a COVID-19-pericyte hypothesis, schematically illustrated in Fig 6. This hypothesis has two key components: 1) The specific organotypic expression of ACE2 together with sufficient S priming proteases in microvascular mural cells should make these cells susceptible for SARS-CoV-2 infection, but their location outside of the endothelium suggests that infection can only occur when the endothelial barrier is broken, allowing blood-borne virus to reach the pericytes. 2) Infection of pericytes may render them dysfunctional, leading to activation of pro-inflammatory and pro-thrombotic responses in neighboring endothelial cells. Immune-attack^62^ on the infected pericytes would cause microvascular inflammation, which may exacerbate leakage and pro-inflammatory and pro-thrombotic responses by the endothelium. Increased vascular leakage would allow more virus to extravasate, infect additional pericytes and intensify and propagate microvascular inflammation and thrombosis. Below, we elaborate the COVID-19-pericyte hypothesis step-by-step, the evidence supporting it, as well as remaining gaps. Finally, we discuss implications of the hypothesis for COVID-19 pathogenesis and therapy.

**Figure 6.**
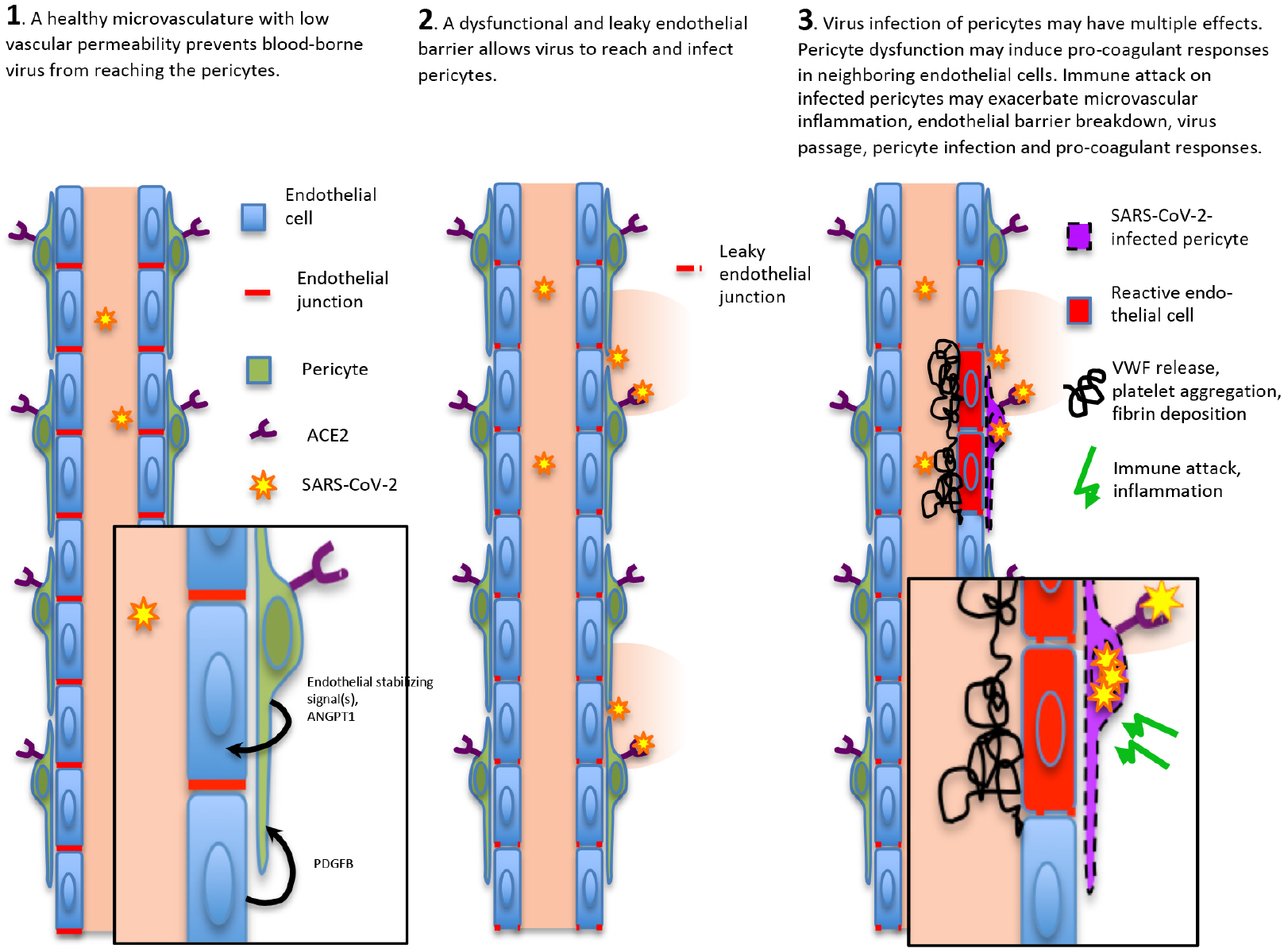
The COVID-19-pericyte hypothesis. The schematic illustration of the COVID-19-pericyte hypothesis depicts a healthy vasculature with low risk of pericyte infection (1), a vasculature in an individual in a COVID-19 risk group with a disrupted endothelial barrier (2); and a SARS-CoV-2 infection of pericytes leading to enhanced expression of Von Willebrand Factor (VWF), platelet aggregation, fibrin deposition and eventually vascular inflammation (3). In the normal, healthy vasculature (1), the intact endothelial barrier precludes SARS-CoV-2 virus to get in contact with the Ace2-expressing pericytes localized behind the endothelial barrier. In individuals with a leaky barrier (2), SARS-CoV-2 can infect pericytes, which in turn causes neighboring endothelial cells to upregulate VWF production, eventually leading to the vascular symptoms observed in many severely ill COVID-19 patients (3).

### ACE2 is expressed by pericytes but not endothelial cells or other perivascular cell types

Accumulating data from analysis of single-cell or single-nucleus transcriptomes suggest that ACE2 is expressed by several epithelial subtypes in the lung, nasal cavity, cornea and conjunctiva and gastrointestinal (GI) tract^7–9^, arguing that one or more of these surface epithelia serve as the entry points for SARS-CoV-2 virus infection. Infection of airway and GI-tract epithelium indeed offers plausible explanation for some of the early and common symptoms of COVID-19 infection, e.g. cough, loss of smell, pneumonia and diarrhea. However, they do not adequately explain the subsequent systemic inflammation and CAC. We surmise that SARS-CoV-2 disseminates into the blood from the sites of primary infection. SARS-CoV-2 RNA has indeed been detected in blood^63^, and, relevant for the COVID-19-pericyte hypothesis, detection of SARS-CoV-2 RNA in blood correlates with further clinical severity^64^.

A cornerstone of the COVID-19-pericyte hypothesis is that pericytes (along with venous and arteriolar VSMCs) are the only ACE2-positive cells in the vasculature. It has been reported that endothelial cells express ACE2 protein^24^, but the data did not distinguish between ACE2 in endothelial versus mural cells. Recent scRNAseq studies also report *ACE2* mRNA expression in cells annotated as endothelial cells^8^. In those cases where we have been able to assess the primary data, we have consistently found evidence for pericyte contamination of endothelial cells. We observe this also in the rare *Ace2/ACE2* mRNA-positive endothelial cells present in our own datasets. Mechanical and enzymatic separation of endothelial cells and pericytes for scRNAseq analysis is problematic because the two cell types are firmly adhered through their joint basement membrane and thereby hard separate without compromising cell viability^29^ (Muhl et al, in press). Without optimized cell-dissociation protocols and positive-negative selection, a substantial proportion of heterotypic endothelial cell-pericyte doublets and cellular fragments will inevitably contaminate the sample^29^.

Besides endothelial cells, we also did not find *Ace2/ACE2* expression in tissue macrophages or other hematopoietic cells, which have been reported to express *ACE2*^8,65,66^. A small subgroup of *Ace2*-positive fibroblasts in the brain were most likely pericyte-contaminated, and this may also be the case for the *ACE2*-positive fibroblasts reported by the Human Cell Atlas Consortium^8^. Moreover, our data show that while pericytes in heart and brain express high levels of *Ace2/ACE2* mRNA, pericytes in lung alveoli and ocular muscle do not express *Ace2*. While additional organs need to be investigated for their mural ACE2 expression, we conclude that vascular *Ace2/ACE2* expression occurs primarily in microvascular mural cells, that the pattern is organotypic, and that there is heterogeneity also within organs, exemplified herein by the heart.

If pericytes become SARS-CoV-2 infected in COVID-19 patients remains to be demonstrated. Pericytes reside within the microvascular basement membrane outside of the microvascular endothelial barrier, and they are therefore normally not in direct contact with blood^50^. The virus particles would reach pericytes only by passing of the endothelial barrier, which is formed by tight and adherens junctions connecting the endothelial cells. In principle, macromolecular passage through the endothelial layer occurs via paracellular paths (where endothelial junctions are discontinuous), transcellular routes (such as endothelial fenestrations) or vesicular transcytosis^67^. However, SARS-CoV2 transport across a healthy endothelial barrier by any of these routes appears unlikely considering the absence of ACE2 in endothelial cells. Given the reported diameter range of 70-150 nm for both SARS-CoV-2 and SARS-CoV-1 (https://www.flickr.com/photos/niaid/albums/72157712914621487)^25,68,69^, the virus should not be able to pass passively through the pores of a healthy endothelium. The physiological upper limit pore size is ≈ 5-12 nm in most types of capillaries including non-sinusoidal fenestrated blood capillaries with diaphragm-containing fenestrae^70^. For brain capillaries with blood-brain barrier, the upper pore size limit is <1 nm. Although the diameter of endothelial fenestrae is comparable to that of the coronavirus particle (i.e. ≈ 100 nm)^71^, the physiological upper pore size limit is ≈ 15 nm in kidney glomeruli and ≈ 60 nm in bone marrow sinusoids, lymph nodes and liver^70,72,73^. Furthermore, even if the virus were able to pass through endothelial pores, it would face a basement membrane with a pore size smaller than the virus diameter^74^. For viral infection of renal proximal tubular epithelial cells, which express ACE2^75^, and for SARS-CoV-2 dissemination into urine^76^, it is therefore conceivable that the virus must translocate through a damaged endothelium located either in the glomerular, or in the peritubular capillaries. In the context of risk factors for severe COVID-19, it is noteworthy that both of these capillary beds are dysfunctional in diabetes^77,78^.

While it appears unlikely that SARS-CoV-2 reaches pericytes in individuals with healthy microvasculature, the situation is different when the endothelial barrier is pathologically disrupted. Breakdown of endothelial junctions and transcytosis has been described in numerous inflammatory conditions^79^, and increased vascular leakage and resulting edema are hallmarks of inflammation. Low-level tissue inflammation, endothelial dysfunction and increased vascular permeability and leakage have been reported in hypertension, diabetes, obesity and aging, and locally in ischemic heart and brain disease, cancer and neurodegenerative diseases^80–88^. All these conditions are risk factors and co-morbidities of severe COVID-19 disease.

An important question that we still lack an answer to is whether severely ill COVID-19 patients have SARS-CoV-2 infected pericytes. Expression of ACE2 with either TMPRSS2 or CTSB/L is sufficient to allow S protein processing and viral entry into cultured cells^2^ and because pericytes express ACE2 and CTSB/L it is plausible that pericytes can become infected if they come in direct contact with virus. Nevertheless, this remains to be demonstrated experimentally or by examination of tissue samples from patients. A recent study showed that SARS-CoV-2 can infect and multiply in human microvascular organoids composed of both endothelial and mural cells^89^, but the exact cellular tropism of the virus in this model remains to be established.

### Pathological consequences of pericyte infection by SARS-CoV-2

While we still lack evidence that pericytes can become infected by SARS-CoV-2 *in vitro* and *in vivo*, we have provided evidence that pericytes regulate pro-thrombogenic responses in the microvascular endothelium. It has previously been shown in mouse models with pericyte hypoplasia that pericyte function is critical for endothelial quiescence, barrier integrity and inhibition of leukocyte adhesion^57–60,90,91^. Herein, we report that endothelial cells in pericyte-deficient mouse microvasculature increase VWF production and release, which would be predicted to promote platelet aggregation and blood coagulation, something we also observe. SARS-CoV-2 infection of pericytes may similarly cause upregulation of VWF production and release in the neighboring capillary endothelial cells. Indeed, increased levels of VWF protein, activity and coagulation factor VIII (which is stabilized by VWF) have been demonstrated in COVID-19 patients with severe disease^11^.

In a recent analysis of the endothelial reaction to pericyte loss, we also found upregulation of several other pro-inflammatory mediators, including angiopoietin-2 (Mäe et al, submitted). However, such pro-inflammatory endothelial changes appear to occur in the absence of severe vascular inflammation. If pericytes become infected by SARS-CoV-2, it is conceivable that they become targets of immune attack^62^, turning a pro-inflammatory endothelial response into fulminant inflammation. Propagation of microvascular inflammation might cause the surge in inflammatory mediators observed in severe COVID-19 cases^92^, including those in the Stockholm region intensive care cohort reported here (Extended Data Table 1). Because the COVID-19-pericyte hypothesis infers that an inflammatory condition, irrespective of cause, would trigger SARS-CoV-2 extravasation, also normally mild inflammation associated with for example mild irritations of the skin or local trauma may be enhanced in SARS-CoV-2 carriers^93^.

We posit the COVID-19-pericyte hypothesis on the basis of a combination of clinical data, single-cell transcriptomics and observations in pericyte-deficient mice. The hypothesis is testable, and will hopefully inform and stimulate additional work aiming at filling the gaps in our data that eventually proves or disproves our hypothesis. The currently missing evidence for pericyte infection by SARS-CoV-2 is an obvious gap that would likely require engagement from the broad research community to be unequivocally filled. A deeper analysis of the range of thrombogenicity and hypercoagulation states in severely ill COVID-19 patients is also warranted, as are detailed morphological analysis of the microvascular morphology with special emphasis on inflammatory hallmarks in multiple organs of deceased COVID-19 patients.

The COVID-19-pericyte hypothesis calls for expanded discussions on the most suitable anticoagulant therapy against CAC^47^. Moreover, the hypothesis calls for introduction of much sought-after preventive therapy among at-risk patients (considering both premorbid risk factors and risk environments). The hypothesis has bearing also on the considerations of type of anti-viral therapy and its modality. As the development of vaccines is sometimes a protracted and uncertain process, systemic delivery of virus-trapping molecules, such as a soluble ACE2 protein^89^, stand out as particularly attractive in a scenario where virus spread through the blood is a disease culprit.

## Methods

### Single cell RNAseq data analysis

#### Mouse heart single cell data integration

ScRNAseq data were obtained from internal mouse heart single cell projects and the published Tabula Muris heart dataset^49^, collectively including diverse cell types in the heart. All samples were obtained from 6-20 weeks old C57Bl6 mice. FACS-based cell capture into 384-well plates with subsequent scRNAseq data generation was conducted using the SMART-Seq2 protocol^94^ and by microfluidic-droplet-based capture by the 10X Genomics protocol. Data processing and clustering were performed using the Seurat package (v. 3.1.1). Cells containing less than 200 expressed genes were filtered out. For the SMART-Seq2 data, cells that generated less than 50,000 reads were filtered out; for the droplet platform, cells containing less than 1000 UMIs were filtered out. Furthermore, genes that were expressed by less than three cells in a dataset were removed. After removing low quality cells from the dataset, the data were normalized using the LogNormalize function, by which feature counts for each cell are divided by the total counts for that cell and multiplied by a scale factor (1 million) and then logarithmically transformed. For integration of different datasets, the integration workflow “Reciprocal PCA” in the Seurat package was implemented, which integrated overall datasets using the mutual nearest neighbor (MNN) cell pairs that shared a common set of molecular features in their PCA spaces. After integration, we obtained a total of 18,378 genes and 10,101 cells for downstream analysis. The function “FindClusters” in the Seurat package was used to identify the clusters with a resolution parameter of 0.5.

#### Mouse lung single cell data integration

The mouse lung datasets were obtained from internal lung single cell projects and the published Tabula Muris lung resource^49^. All samples were from 10-19 weeks old C57Bl6 mice. Data integration and clustering analysis for the lung were performed with the same methods as for the mouse heart data described above. We obtained a total of 20,114 genes and 11,085 cells in the integrated lung dataset.

#### Mouse brain single cell data integration

Mouse brain datasets were integrated from two internal brain single-cell projects and one published (the Tabula Muris brain resource)^49^. The internal datasets included one unpublished and one previously published brain vasculature dataset^30^. The cells were from 10-19 weeks old C57Bl6 mice. Data integration and clustering analysis were performed with the same methods as for the mouse heart and lung data described above. We obtained a total of 12,940 cells and 17,779 genes in the integrated brain dataset.

#### Bar plot visualization of integrated datasets

In order to provide detailed visualizations of the primary gene expression data cell-by-cell for each cluster of the integrated dataset, we created bar plots using the normalized counts from each cell. In these graphs, a bar represents a cell and is colored according to its data source. The data source abbreviations “ss2” and “droplet” in the legend represent the SMART-Seq2 protocol and microfluidicdroplet-based capture by the 10X Genomics protocol, respectively. For the integrated dataset of lung, “TJA_ss2”, “CBZ_ss2” and “H_droplet” represent in-house unpublished datasets; “TM_ss2” and “TM_droplet” represent published Tabula Muris data^49^. For the integrated dataset of heart, “L_ss2”, “S_ss2” and “H_droplet” represent in-house unpublished datasets; “TM_ss2” and “TM_droplet” represent published Tabula Muris data. For the integrated dataset of brain, “C_ss2” and “K_droplet” represent in-house unpublished datasets; “TM_ss2” represents published Tabula Muris data.

#### Human heart

The human heart single cell data were extracted from a published study^54^, and only cells from healthy donors were included in the current analysis. The Seurat package (version 3.1.1) was used for raw data processing, filtering, normalization, clustering and further downstream analysis^95^. Cells that had less than 500 expressed genes were filtered out. Genes expressed in less than 10 cells were also filtered out. In total, 8383 single cells from 14 previously healthy organ donors (12 males and 2 females) qualified for downstream analysis. The gene expression levels in each cell were normalized to a total read counts of 100,000 per cell. The top 2,000 variable genes in the dataset were used for linear dimensional reduction of the data using the PCA method. The first 30 principal components were used for UMAP visualization and clustering of the cells using default parameters in Seurat pipeline.

#### Human lung

The human lung single cell data were obtained from a published study^56^. In total, there were 76,070 qualified single cells from 16 individuals (8 donors and 8 lung fibrosis patients). The Seurat package (v3.1.1) was used for data processing. The cell type annotations shown are the same as in the original paper^56^. To generate the UMAP layout of the data, the reciprocal PCA method in the Seurat integration pipeline was used.

#### Identification of pericyte contamination of other cell types

To identify pericyte contamination in other cell types, including endothelial cells, fibroblast-like cells and cardiomyocytes, we examined the expression of several previously well-characterized pericyte-specific markers, including *Kcnj8, Pdgfrb* and *Abcc9*. Their expression profiles in *Ace2*-positive and *Ace2*-negative cells were compared in a random selection of equal numbers of cells, and the heat map results were visualized with pheatmap package (version 1.0.12) in R software.

### Mice

The following mouse strains were used: *Pdgfb*^ret^ (*Pdgfb*-tm(ret))^61^, *Cspg4*-DsRed (The Jackson Laboratory, Tg(Cspg4-DsRed.T1)1Akik/J, *Pdgfrb-GFP* (Gensat.org. Tg(Pdgfrb-eGFP) JN169Gsat/Mmucd)^28^, *Cldn5-GFP* (Tg(Cldn5-GFP)Cbet/U), *Acta2^GFP^* (The Jackson Laboratory, Tg(*Acta2*-GFP)1Pfk), *Prox1-GFP* (Tg(Prox1-EGFP)KY221Gsat/Mmcd). All mice were backcrossed on a C57BL6/J genetic background. Adult mice, about 3 months of age and of both sexes, were used for experiments. Animal protocols were approved by either the Uppsala Ethical Committee on Animal Research (permit numbers C224/12, C115/15, C111515/16), or by the Stockholm/Linköping Ethical Committee on Animal Research (permit ID 729). All animal experiments were carried out in accordance with their guidelines.

### Immunofluorescence staining

#### *Pdgfb*^ret/+^ and *Pdgfb*^ret/ret^ *brains*

Mice under full anesthesia were euthanized by transcardial perfusion with Hanks balanced salt solution (HBSS, cat. #14025092, GIBCO) followed by 4% buffered formaldehyde (cat. #02178, Histolab). Brains were removed and post-fixed in 4% buffered formaldehyde for 4 h at 4 °C. Sagittal and coronal vibratome sections (50-75 μm) were incubated in blocking/permeabilization solution (1% bovine serum albumin, 2.5% donkey serum, 0.5% Triton X-100 in PBS) overnight at 4 °C, followed by incubation in primary antibody solution for two nights at 4 °C, and subsequently in secondary antibody (Jackson ImmunoResearch and Invitrogen) solution, overnight at 4 °C. A list of the used primary antibodies is presented in Extended Dataementary Table 1. Sections were mounted in ProLong Gold Antifade mounting medium (cat. #P36930, Life Technologies). Micrographs were taken with a Leica TCS SP8 confocal microscope (Leica Microsystems). All confocal images are represented as maximum intensity projections and were adjusted for brightness and contrast using Fiji v1.52p and Adobe Photoshop CC 2019.

#### Blood-brain barrier permeability

For blood-brain barrier integrity assessment, dextran (100 μg/g body weight) conjugated to tetramethylrhodamine (cat. #D1818, Life Technologies) was injected intravenously into the tail vein 16 hours before sacrifice, respectively^57^. For tracer *in situ* analysis, anaesthetized animals were perfused transcardially for 5 min with HBSS followed by 4 min with 4% buffered formaldehyde. Where after the brains were processed as described under *Immunofluorescence staining*.

#### Cryo-sections from brain, heart and lung

Tissues were harvested from euthanized mice without perfusion and fixed by immersion in 4% formaldehyde for 4-12h at 4 °C, followed by immersion in 20% sucrose/PBS solution for at least 24h at 4°C. Thereafter, tissues were embedded for cryo-sectioning and sectioned on a CryoStat NX70 (ThermoFisher Scientific) to 14 or 30 μm thick sections collected on SuperFrost Plus glass slides (Metzler Gläser). Sections were allowed to thaw at RT and thereafter blocked for > 60 min at RT with blocking-buffer (serum-free protein blocking solution, DAKO), supplemented with 0.2% Triton X-100 (Sigma Aldrich), followed by sequentially incubation with primary antibodies (overnight at 4 °C) (Extended Data Table 1) and corresponding fluorescently conjugated secondary antibodies (1h at RT) together with 10 μg/ml *Hoechst 33342* (trihydrochloride, trihydrate, ThermoFisher Scientific). Sections were mounted with ProLong Gold Antifade mounting medium, and micrographs acquired and graphically handled as described above.

## Legends to Extended Data Tables and Figures and Videos

**Extended Data Table 1: Clinical, inflammatory and coagulation parameters in 20 ICU-admitted Covid-19 patients.**

The ICU patients had a median age of 63 years (range 40-73), the majority being male (90%) and having one or more risk factors prior to admission (85%). The length of ICU stay and duration of invasive mechanical ventilation reveal the severity of disease and treatment challenge. Most of them required mechanical ventilation in prone position due to critical hypoxia with periods of high-inspired fractions of oxygen (FiO_2_) and a low arterial oxygen tension to FiO_2_ ratio index (PFI). Continuous parameters are depicted as median (range) and categorical parameters as number (%). Intensive Care Unit (ICU), ratio arterial oxygen tension (kPa)/ fraction inspired oxygen (FiO_2_) (PFI), C-reactive protein (CRP), procalcitonin (PCT), interleukin-6 (IL-6), interleukin-1 (IL-1), tumour necrosis factor alpha (TNF-alpha), internationalized normalized ratio (INR), activated partial thromboplastin time (APTT). Risk factors include renal disease, hypertension, cardiovascular disease, diabetes mellitus, chronic pulmonary disease and immunosuppressive therapy^15,41,42^.

**Extended Data Figure 1. *Ace2* mRNA in brain cells other than mural cells is due to pericyte-contamination**

***a*.** Top enriched transcripts (red arrow indicates *Ace2* at 15^th^ place) in pericytes and venous SMC as compared to other vascular and perivascular cell types deduced from the brain vascular scRNAseq database at ***b*.** Bar plot excerpt from http://betsholtzlab.org/VascularSingleCells/database.html showing the expression of *Ace2* in different vascular and perivascular cell types. ***c*.** Magnified view of indicated part of ***a*** comparing the expression of *Ace2, Cnn1* and *Kncj8*. The abbreviations are as follows: PC, Pericytes; SMC, Smooth muscle cells; MG, Microglia; FB, Vascular fibroblast-like cells; OL, Oligodendrocytes; EC, Endothelial cells; AC, Astrocytes; v, venous; capil, capillary; a, arterial; aa, arteriolar. ***d*.** Excerpts from http://betsholtzlab.org/VascularSingleCells/database.html showing bar plots across VSMC subtypes of arterial VSMC-specific genes that anti-correlate with *Ace2*. Red arrows point at the *Ace2*-negative part of the cluster. Cell type abbreviations: SMC, Smooth muscle cells; v, venous; a, arterial; aa, arteriolar. ***e*.** Expression of selected known brain pericyte specific markers in 23 *Ace2*-positive endothelial cells (top) and 23 randomly selected *Ace2*-negative endothelial cells (bottom). Colors indicate pericyte marker as shown, and the frequency of their expression is provided in the tables. ***f*.** Heat map display of pericyte and fibroblast markers in *Ace2*-positive (*Ace2*+) and *Ace2*-negative (*Ace2*-) FB2 fibroblasts and enriched genes in the *Ace2+* cells. ***g*.** Heat map display of brain cell type specific markers of different vascular and perivascular cells in *Ace2+* and randomly selected equal number of *Ace2*-endothelial cells. Only pericyte markers are enriched in *Ace2+* endothelial cells, not arterial VSMCs or astrocyte markers. ***h*.** Heat map display of the 50 top enriched genes in the three *Ace2+* cells (black arrowheads) without pericyte marker expression does not reveal a statistically significant commonality between these cells. Most of the genes listed in **d** are highly expressed in neurons, suggesting an RNA smearing artifact.

**Extended Data Figure 2. ACE2 protein expression in the adult mouse brain cortex and spinal cord.**

***a-c*.** Confocal microscopy images of sections from brain cortex (***a-c***) and spinal cord (***d-f***) IF stained using antibodies and *Pdgfrb-GFP* marker as indicated. Horizontal image pairs depict the same microscopic field with different label. Note the strong ACE2 staining of capillary pericytes and the co-expression with CD13 (***a***) and *Pdgfrb*-GFP (***b***). Mural cells in terminal arterioles are weakly ACE2 positive (***c*,** arrowheads and ***e***, arrow), whereas mural cells of larger arterioles are ACE2 negative (arrows in ***c*** and ***f***).

**Extended Data Figure 3. ACE2 protein expression in the adult mouse eye.**

Confocal microscopy images of cross-sections of adult mouse eyes. Each row of four images shows the same microscopic field with different combinations of label, as indicated. Double insets display the same field with different label as indicated, aiming to illustrate: (***b***) that *Pdgfrb*-GFP-positive pericytes in ocular skeletal muscles are ACE2-negative; (***d,l***) that capillary pericytes in the choriocapillaris are ACE2-positive but the VSMC of its feeding arteries are ACE2-negative; that the feeding arterioles of the ciliary body vasculature are ACE2-negative (***g***) but the ciliary body pericytes ACE2-positive (h); that the retinal pericytes and VSMC of terminal arterioles are ACE2-positive but the larger arteries are ACE2-negative (***k***). Asterisks in ***c,g,k,o*,** indicate ACE2-positive retinal pericytes. Arrowheads in ***c,k,s*** point at ACE2-positive pericytes in the choriocapillaris. Arrow in ***t*** points at an ACE2-negative artery in the choriocapillaris.

**Extended Data Figure 4. *ACE2* expression in human brain pericytes.**

***a***. The expression of known pericyte markers in *ACE2+* cells from developing human prefrontal cortex. b. Heat map display of gene expression in two *ACE2+* cells and five endothelial cells retrieved from a human glioblastoma scRNAseq dataset.

**Extended Data Figure 5. *Ace2* expression in mouse heart endothelial cells is due to pericyte contamination.**

***a*.** Heat map overview of the expression of pericyte and endothelial markers in *Ace2+* pericytes and *Ace2+* and *Ace2*-endothelial cells in the adult mouse heart. Arrows indicate the position of three *Ace2*+ cells without other known pericyte markers. ***b***. The expression of known pericyte-specific markers in the 21 *Ace2*-positive endothelial cells (top) and 21 randomly selected *Ace2*-negative endothelial cells (bottom). The expression of markers in the same cell is piled up. ***c*.** Heat map overview of the top enriched genes in the three *Ace2*-positive cells (arrow highlighted) without other known pericyte marker expression. There was no apparent common gene profile among the three cells, suggesting that they do not represent a specific cell type or cell state.

**Extended Data Fig 6: ACE2 expression in pericytes in the adult human heart.**

**a.** UMAP visualization of human heart cells. Colors indicate cluster assignment. Annotations were based on the expression of canonical markers for each indicated cell type. Red rrow indicates the pericyte cluster. **b**. Bar plots of the normalized expression levels of *ACE2* and the indicated cell type markers in the same clusters as displayed in the UMAP plot. Each bar represents a cell. The pericyte cluster (#4) is denoted by a red arrow. The endothelial cluster (#2) is denoted by a black arrow. **c**. The expression of known pericyte markers in 70 *ACE2*-positive endothelial cells (top) and 70 *ACE2*-negative endothelial cells (bottom). Each bar is a cell and marker expression is piled up. For each group, statistics of pericyte (PC) markers is summarized in the corresponding table, including numbers and percentages of cells with no pericyte marker expressed, one or more pericyte marker, and two or more pericyte markers (indicated by # and *), respectively.

**Extended Data Figure 7. ACE2 protein expression in the adult mouse lung.**

Confocal microscopy images of sections from adult mouse lung stained with the indicated antibodies (ACE2, CD31 for endothelium, NG2 for mural cells, CD68 for macrophages) and the transgenic reporter *Pdgfrb*-GFP for mural cells. Each row of images depicts the same microscopic field with different labels. ACE2 IF signal is only detected in bronchial epithelium (strong) and AT-2 cells (weak). Note the absence of ACE2 IF signal in endothelial cells, in mural cells in the alveolar region and in CD68 positive macrophages. Double insets depict the same region, one with multiple IF labels and one with ACE2 only. Arrows point at rare ACE2:*Pdgfrb*-GFP double positive mural cells located close to large bronchi. Bottom panels show tracheal capillaries with attached ACE2-positive pericytes. Arrowheads in the magnified inset point at four pericyte somata. Scale bars: 100 μm.

**Extended Data Figure 8. *ACE2* expression in the human lung.**

**a.** UMAP visualization of human lung scRNAseq data (left) with cluster annotations taken from the original paper and its marker basis in the text box (top 10 markers for each cell type)^56^. *ACE2* expression superimposed onto the UMAP image (right, dark color represents higher expression and grey color represents *Ace2*-cells). **b.** Bar plot display of *ACE2* mRNA shows that it is localized to the clusters annotated as AT-2 (*SFTPC*+), basal (*KRT5*+), multiciliated (*FOXJ1*+) and club (*SCGB1A*+) cells.

**Extended Data Figure 9. Co-expression analysis of *Ace2, Tmrpss2, Ctsl and Ctsb* in mouse lung, brain and heart.**

Bar plots overview of *Ace2, Tmrpss2, Ctsl and Ctsb* expression across the scRNAseq meta-analysis datasets of lung, brain and heart. Each bar represents a single cell and is colored according to the indicated data source (see Materials and Methods). Cluster annotations have been described in previous figures. Arrows point at the Ace2+ clusters, which in lung represent AT-2 (left) and multiciliated cells (right), in brain: from left to right, pericytes, arteriolar VSMCs, venous VSMCs and pericyte-contaminated endothelial cells, and in heart: pericytes.

**Extended Data Figure 10. Pericyte loss in *Pdgfb^ret/ret^* heart.**

Confocal microscopy images of sections from adult mouse *Pdgfb^ret/+^* and *Pdgfb^ret/ret^* hearts stained with the indicated antibodies, CD31 for endothelial cells and NG2 for mural cells. Areas were selected from the left ventricular wall. Horizontal pairs of images depict the same microscopic field with different labels as indicated. The bottom panels show magnified view of marked insets. Note the reduced density of NG2 positive cells in the *Pdgfb^ret/ret^* microvasculature. Scale bars: 100 μm.

**Extended Data Figure 11. VWF immunofluorescence analysis shows specific localization to Weibel–Palade bodies.** Localization of VWF staining to rod-shaped vesicles (Weibel-Palade bodies) in the endothelial cytoplasm, i.e. underneath the CD31-positive endothelial membrane, which in turn is underneath the ACTA2-positive VSMC coat of a brain arteriole in a control (*Pdgfb*^ret/+^) mouse. This staining served as a control for the specificity of the antibody and quality of the immunofluorescence analysis. See also Extended Data Video 1. Scale bar: 50 μm.

**Extended Data Figure 12. Platelet adhesion and aggregation in *Pdgfb^ret/re^* vessels.**

**a.** Visualization of platelets using anti-CD41 and fibrinogen (FBG) leakage and fibrin deposition in blood vessels (CD31; PECAM1) of *Pdgfb^ret/ret^* mouse brains. Arrows in the top panel point at a fibrin clot. Arrows in the bottom panel highlight platelets and local fibrin deposition. **b.** VWF release into the interstitium coincides with 70 kDa dextran tracer leakage (red) into the brain parenchyma (yellow arrows) and COL4 (COL4A1) visualizes the basement membrane of the vasculature. CD13 (ANPEP) visualizes residual mural cells. Note the anti-correlation between presence of mural cells and VWF expression. Scale bars: 25 μm in **a** and 50 μm in **b.**

**Extended Data Video 1. VWF immunofluorescence analysis shows specific localization to Weibel–Palade bodies.** Z-scan of the arteriole shown in Extended Data Figure 11 provides a clear view of the localization of the VWF-positive vesicles in the endothelial cytoplasm. Video download is available at http://betsholtzlab.org/Publications/2020/Extended_DataVideo1.html

## Acknowledgements

We thank Cecilia Olsson, Pia Peterson, Jana Chmielniakova, Helene Leksell, Sonja Gustafsson, Byambajav Buyandelger and Elisabeth Raschperger for technical help. This study was supported by grants from the Swedish Cancer Society (C.B., U.L.), the Swedish Research Council (C.B., U.L), the Swedish Brain Foundation (C.B., U.L.), the Erling-Persson Family Foundation (C.B., U.L.), Knut and Alice Wallenberg Foundation (C.B., T.M.), The European Union (C.B.), the Leducq Foundation (C.B.), the Louise Jeantet Prize (C.B.), The Anders Jahre Medical Prize (C.B.) and ERC advanced grant (C.B.) and consolidator grant (T.M.). C.B. and E.H. were supported by grants from AstraZeneca through the ICMC. S.S. was supported by a postdoctoral fellowship from the Deutsche Forschungsgemeinschaft (STR 1538/1-1) and a non-stipendiary longterm fellowship from the European Molecular Biology Organization (ALTF 86-2017).

## Author contributions

C.B. conceived the COVID-19-pericyte hypothesis and developed it together with T.A., X-R.P., L.I.E. and U.L. L.H. did the bioinformatics analysis. M.A.M., E.V.L. and K.N performed analyses of *Pdgfb* mutant mice. Y.S. designed the scRNAseq metaanalysis pipeline and applied it together with R.P. M.A.M., L.M. and R.P. performed ACE2 immunofluorescence analysis. M.J.F and A.O. explored and analyzed ICU patient data. L.M., M.A.M., J.L., G.G., L.Z., Y.X., S.L., G.M. S.S., A.O., M.R., A.A., J.B. M.V., K.B., E.H., K.A. and T.M. provided unpublished scRNAseq data. C.B. assembled the data. C.B. and U.L. wrote the manuscript with significant input fro K.A. All authors reviewed and edited the text.

