## Extended Data and Supplement for "Pericyte-specific vascular expression of SARS-CoV-2 receptor ACE2 – implications for microvascular inflammation and hypercoagulopathy in COVID-19"

**Extended Data Table 1. Description of a cohort of 20 ICU patients diagnosed with COVID-19.**

| | | |
|---|---|---|
| ***Patient characteristics in ICU*** | | |
| Age (years) | 63 (40-73) | |
| Sex (male/female) | 18/2 | |
| Patients with riskfactors (n) | 17/20 (85%) | |
| Days with sickness symtoms before ICU admission (days) | 10 (2-21) | |
| Days in ICU (days) | 23 (5-37) | |
| Duration of mechanical ventilation (days) | 22 (4-36) | |
| PFI minimum | 11.8 (6-20) | |
| $FiO_2$ maximum | 0.73 (0.5-1.0) | |
| PEEP maximum | 14 (8-18, 10) | |
| Patients treated in prone position (n) | 17/20 (85%) | |
| Patients treated with CRRT (n) | 10/20 (50%) | |
| Patients with confirmed tromboembolism (n) | 6/20 (30%) | |
| Secondary infection | 12/20 (60%) | |
| Therapeutic anticoagulation (n) | 8/20 (40%) | |
| Profylactic anticoagulation (n) | 15/20 (75%) | |
| Patients with profylactic ASA (n) | 10/20 (50%) | |
| ***Patients Systemic Inflammatory Biochemistry*** | | ***Reference values*** |
| White cell count max | 19.6 (13.6-36.1) x 10 (9)/L | 3.5-8.8 x 10 (9)/L |
| Neutrophile count max | 16.7 (3.1-34.1) x 10 (9)/L | 1.6-5.9 x 10 (9)/L |
| Lymfocyte count min | 0.5 (0.2-7.2) x 10 (9)/L | 1.1-3.5 x 10 (9)/L |
| IL-1 max | 8.3 (5-67) ng/L | <5 ng/L |
| IL-6 max | 670 (83-49544) ng/L | <7 ng/L |
| TNF-alpha max | 36.7 (17.8-67.0) ng/L | <12 ng/L |
| Ferritin max | 2169 (618-9752) µg/L | 10-150 µg/L |
| CRP max (mg/ml) | 388 (256-642) mg/L | <3 mg/L |
| PCT max (µg/L) | 9.2 (1.6-306) µg/L | <0.5 µg/L |
| ***Patients Systemic Coagulation Biochemistry*** | | ***Reference values*** |
| SR | 140 (87-140) mm | <10 mm |
| INR | 1.3 (1.0-8.0) | <1.2 |
| APTT | 34 (25-119) s | 20-30 s |
| TPK min | 199 (19-530) x 10 (9)/L | 145-348 x 10 (9)/L |
| TPK max | 487 (89-1171) x 10 (9)/L | 145-348 x 10 (9)/L |
| D-dimer max | 8 (2-35) mg/L FEU | <0.5 mg/L FEU |
| AT III min | 0.78 (0.58-1.04) kIE/L | 0.8-1.2 kIE/L |
| Fibrinogen max | 9.3 (5.8-15.8) g/L | 2-4.2 g/L |

Data are presented as median (min-max, range) or numbers (percentage)

### Extended Data Fig 1

*a*

Top 20 enriched genes in group (G)1 vs. G2
G1= PC, vSMC
G2= aaSMC, aSMC, MG, FB1, FB2, OL, EC1, EC2, EC3, vEC, capEC, aEC, AC

| # | Symbol | G1_avg. | G2_avg. | Fold(G1/G2) |
|---|---|---|---|---|
| 1 | Sebox | 57.02 | 0.36 | 157.05 |
| 2 | Nts | 39.39 | 0.39 | 100 |
| 3 | Nodal | 104.94 | 1.19 | 88.47 |
| 4 | Iqsec3 | 18 | 0.24 | 76.22 |
| 5 | Atp13a5 | 4185.95 | 102.73 | 40.75 |
| 6 | 5033404E19Rik | 81.32 | 2.02 | 40.32 |
| 7 | Art3 | 811.79 | 22.17 | 36.62 |
| 8 | Nxph4 | 34.04 | 1.22 | 27.93 |
| 9 | Vtn | 15514.65 | 559.46 | 27.73 |
| 10 | Sfrp2 | 49.54 | 1.85 | 26.73 |
| 11 | Arl5c | 14.27 | 0.58 | 24.73 |
| 12 | Slc6a20a | 790.97 | 32.4 | 24.41 |
| 13 | Tmem26 | 10.08 | 0.42 | 24 |
| 14 | Plxdc1 | 338.27 | 14.86 | 22.77 |
| 15 | Ace2 | 403.42 | 18.01 | 22.4 |
| 16 | Abcc9 | 741.44 | 33.8 | 21.93 |
| 17 | Pla1a | 305.17 | 14.18 | 21.53 |
| 18 | P2ry14 | 1222.11 | 57.62 | 21.21 |
| 19 | Kcnj8 | 1105.9 | 52.58 | 21.03 |
| 20 | Uchl1 | 146.45 | 7.17 | 20.42 |

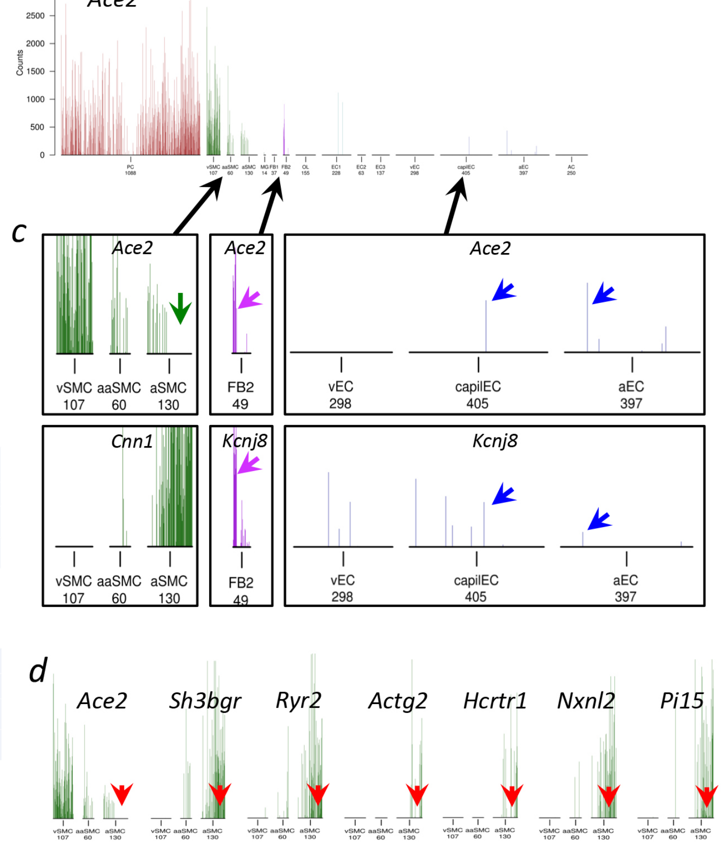


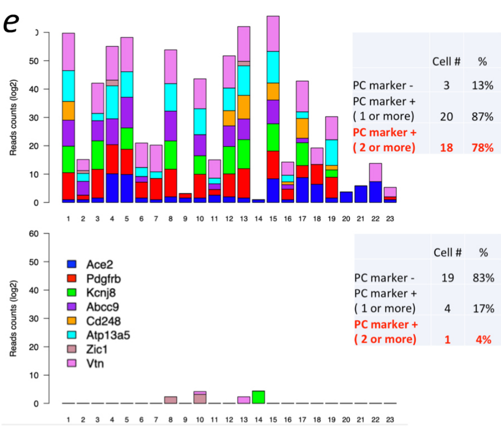


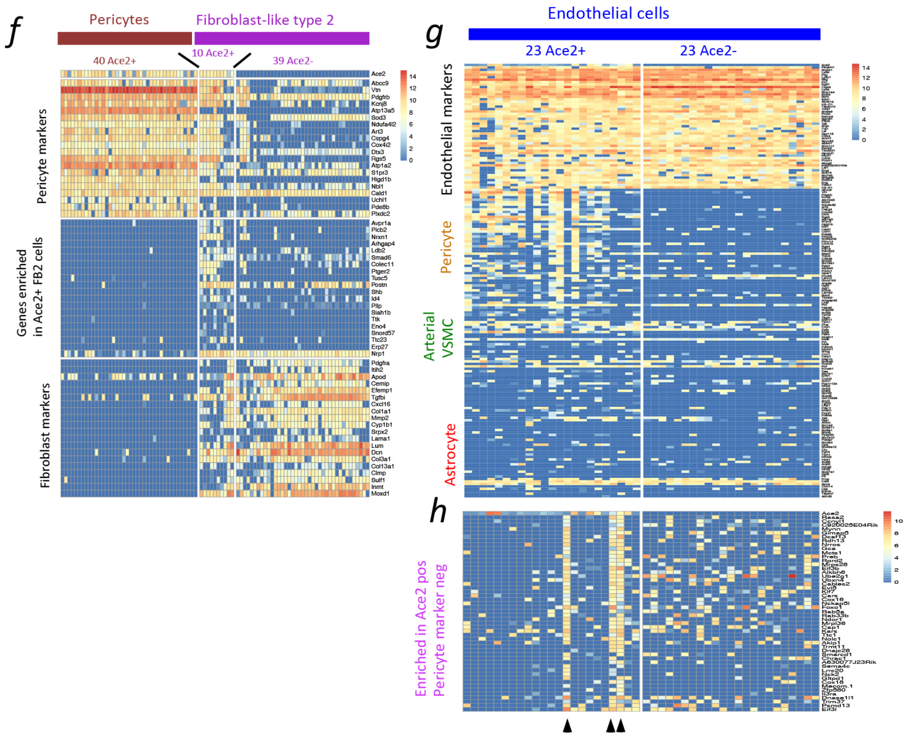

Extended Data Fig 2

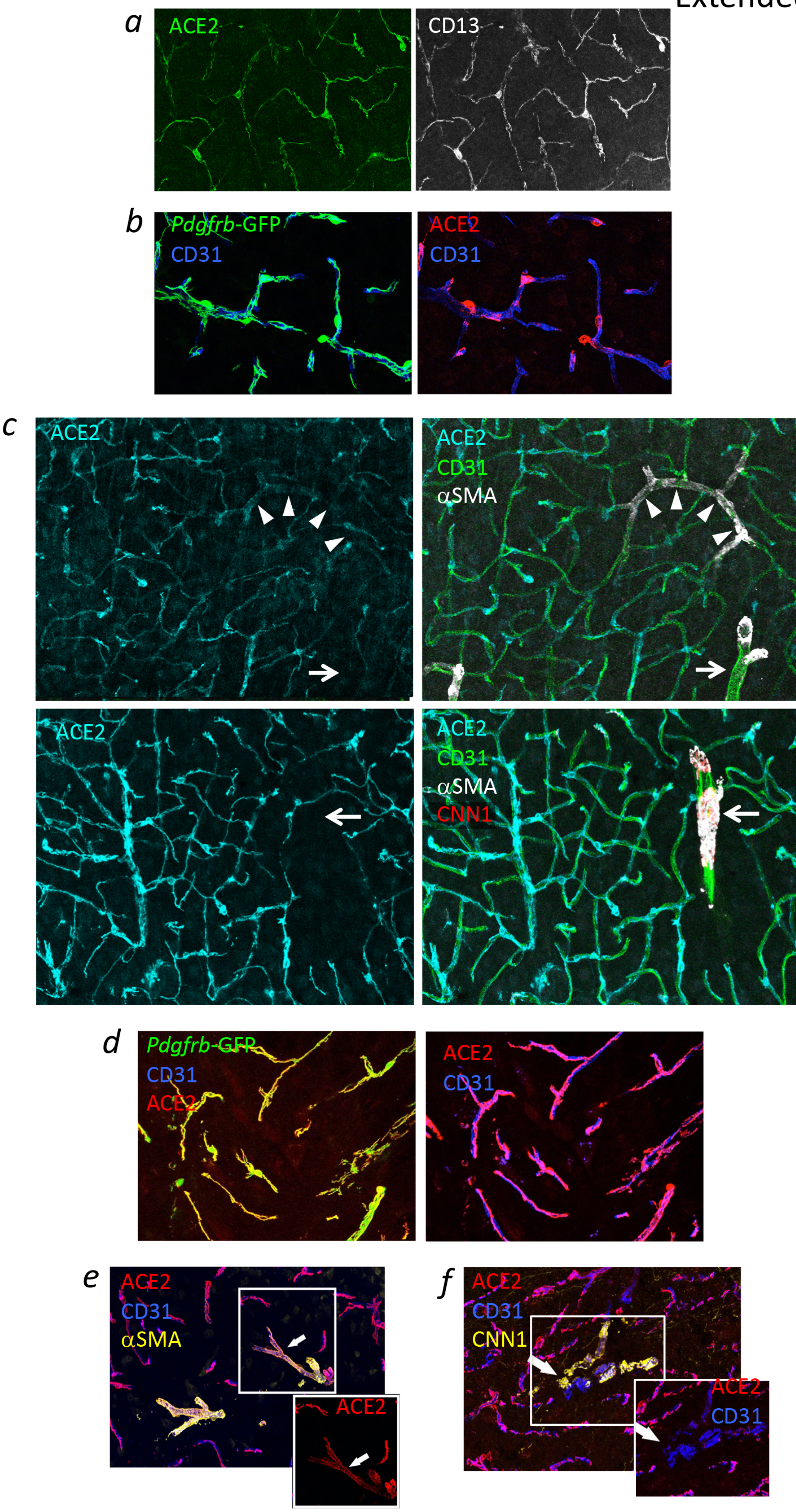

Extended Data Fig 3

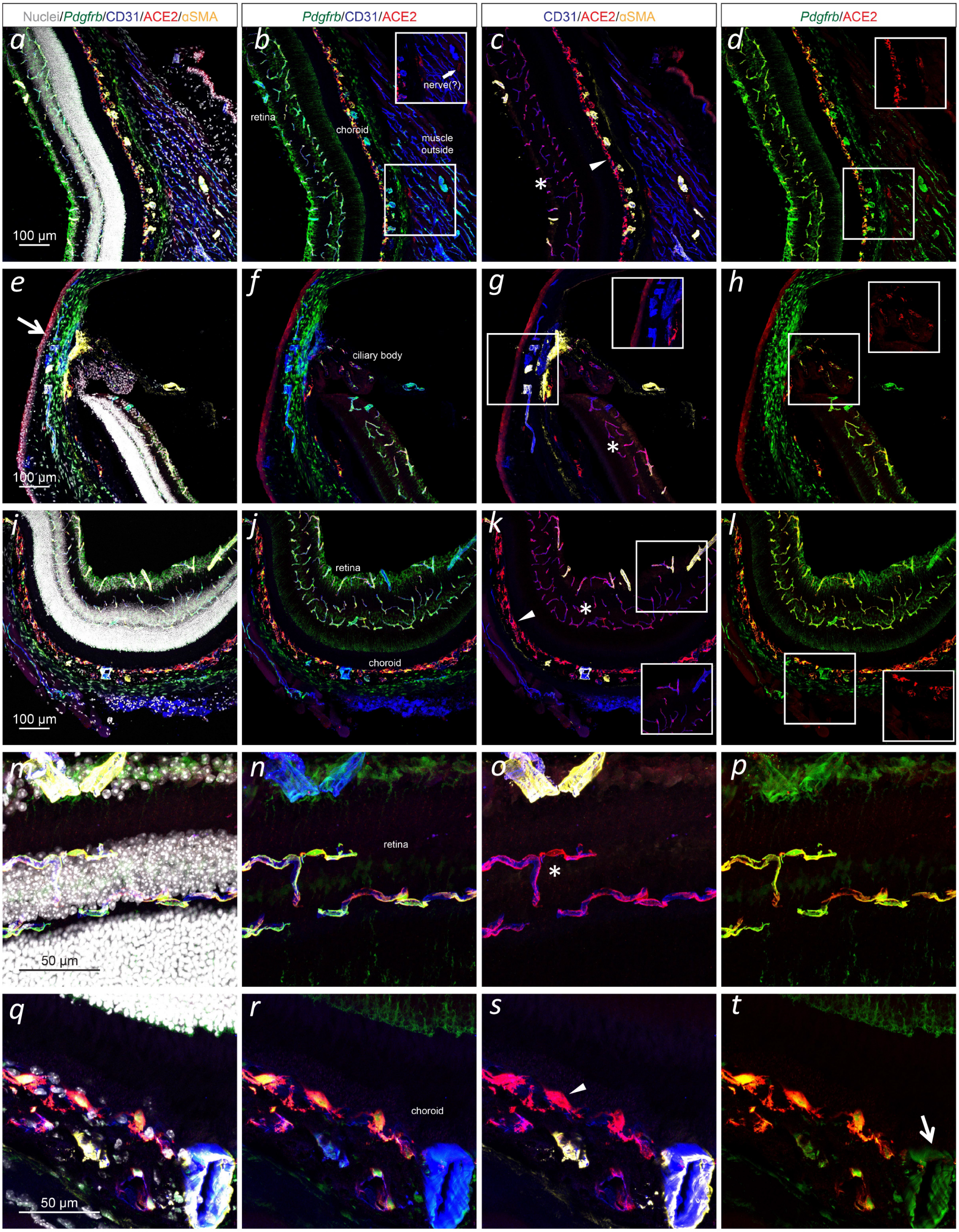

### Extended Data Fig 4

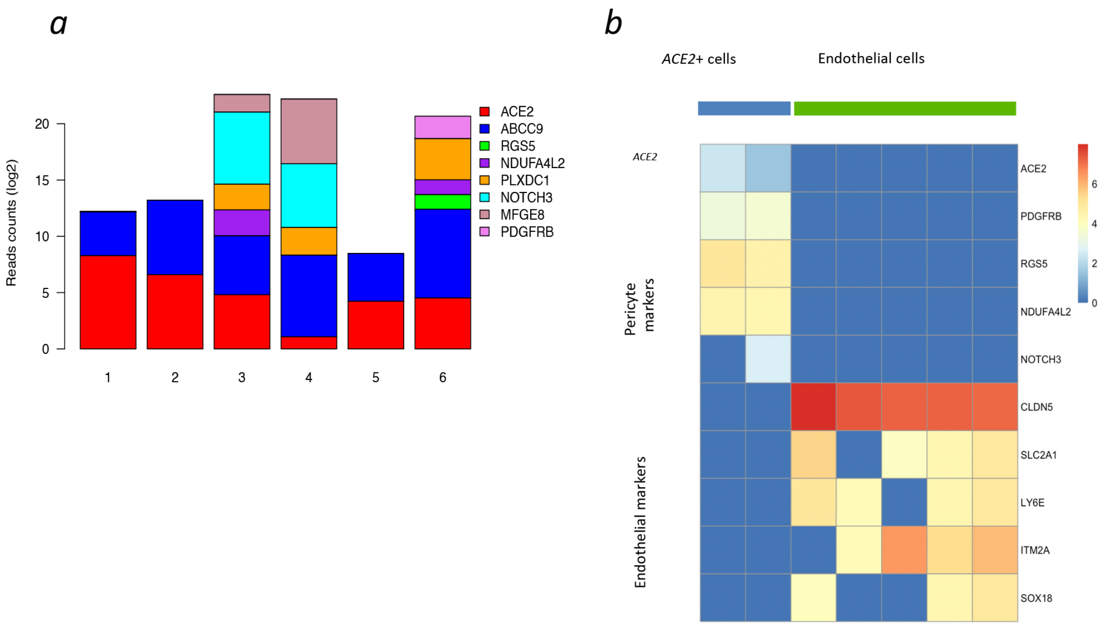

### Extended Data Fig 5

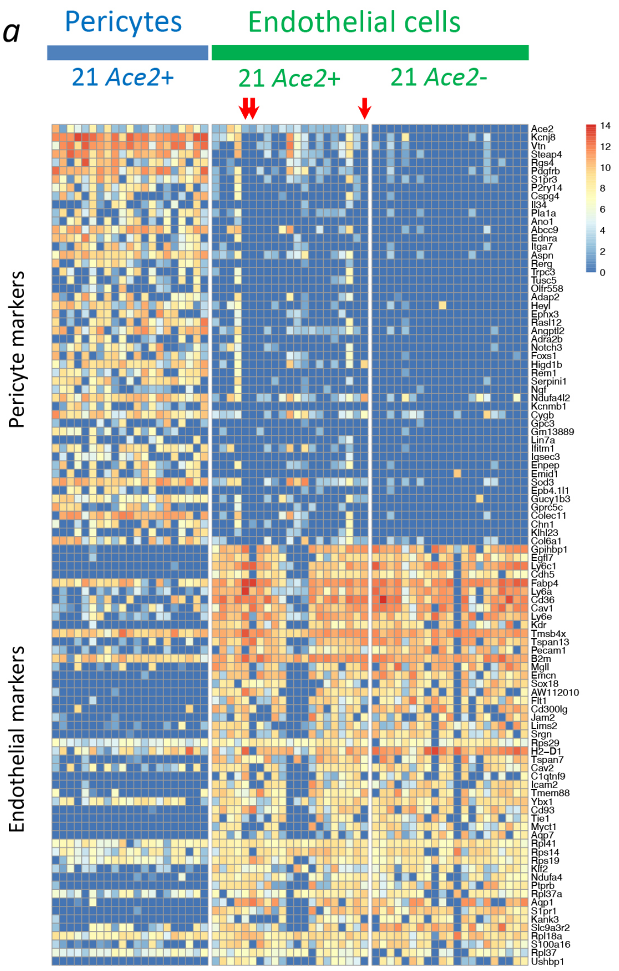


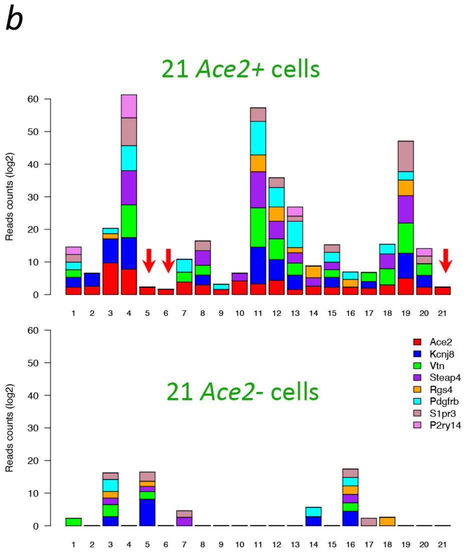


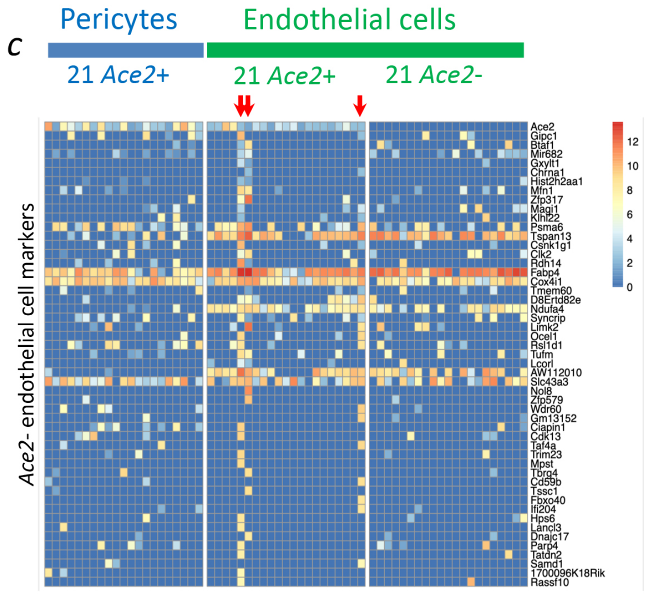

### Extended Data Fig 6

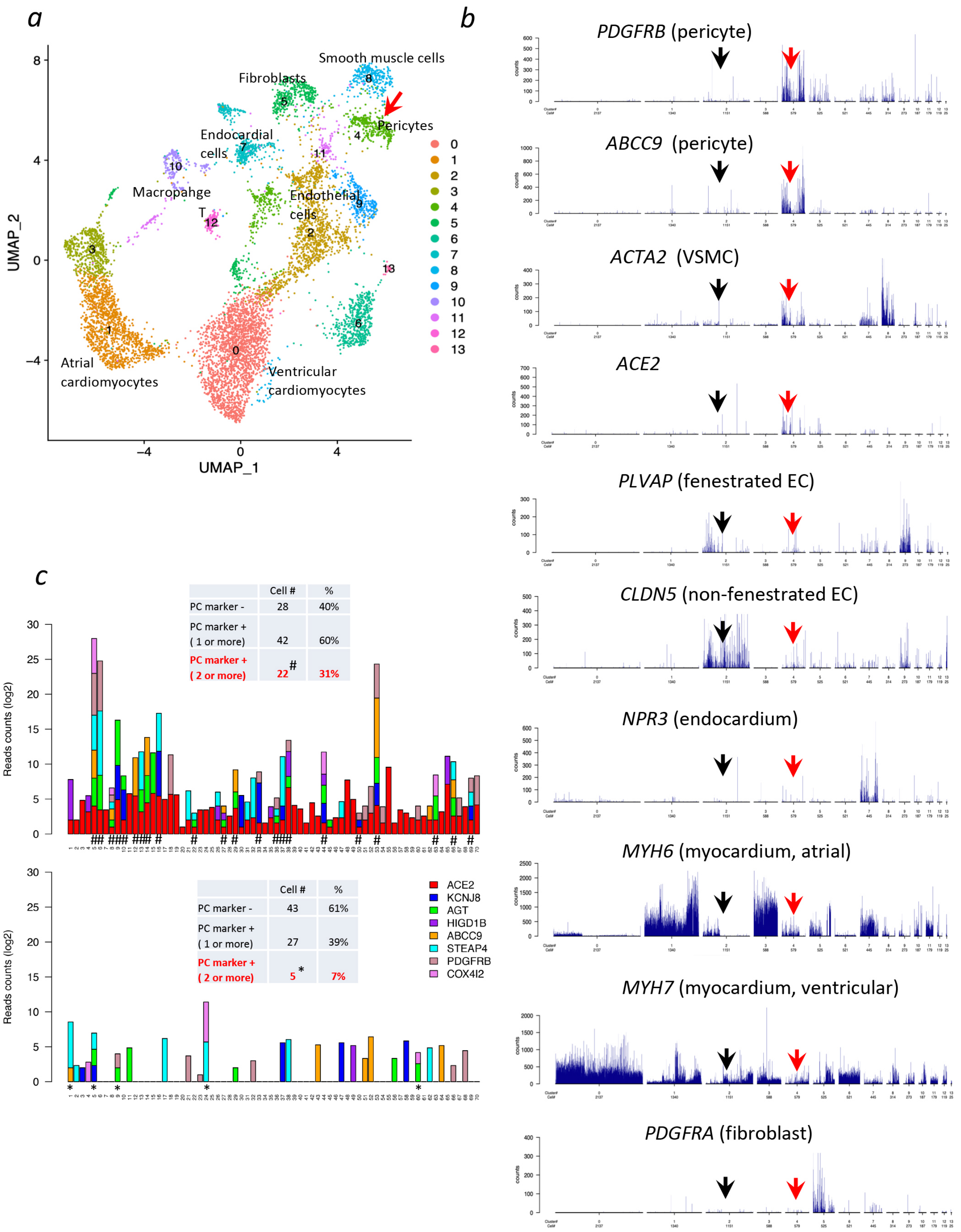

Extended Data Fig 7

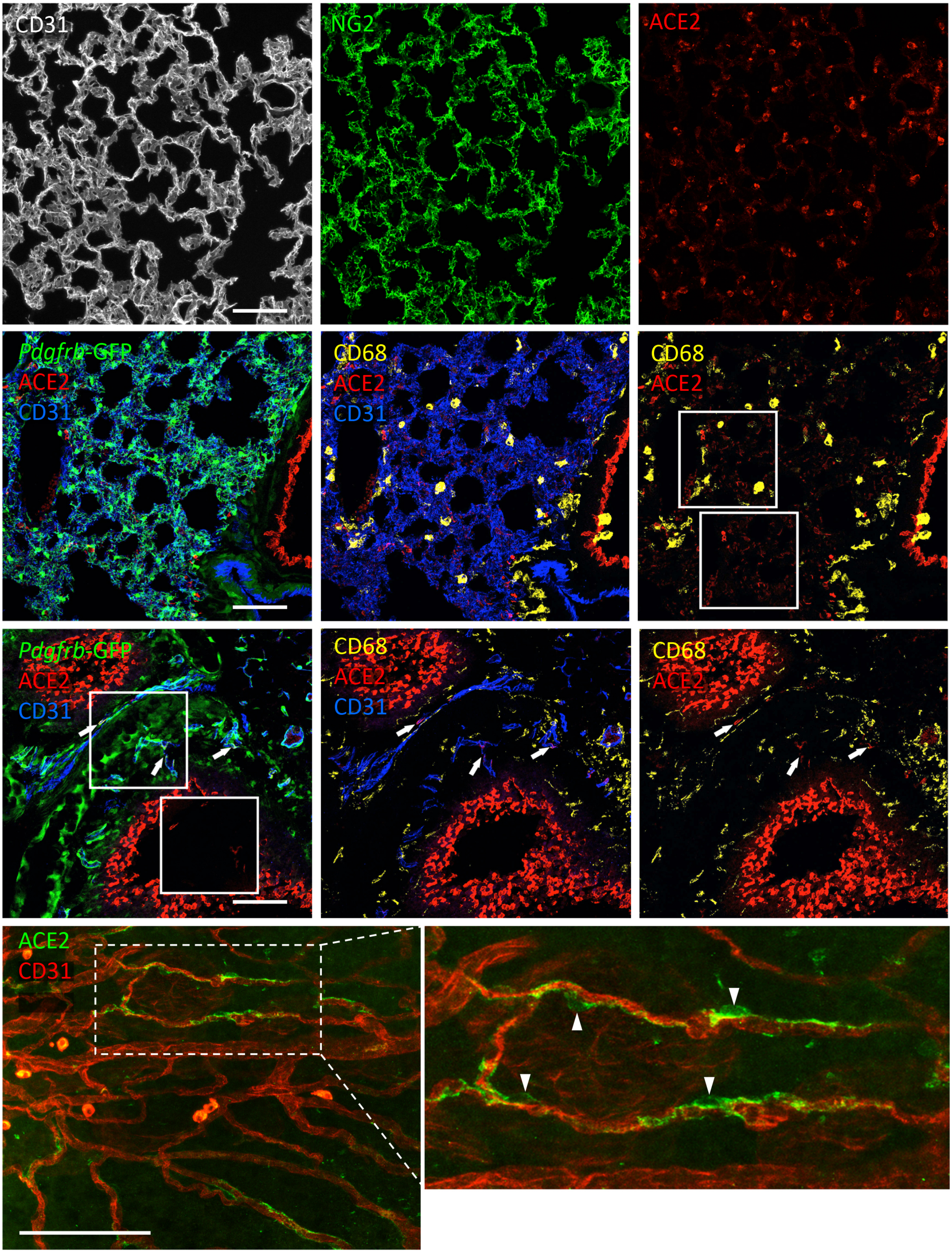

Extended Data Fig 8

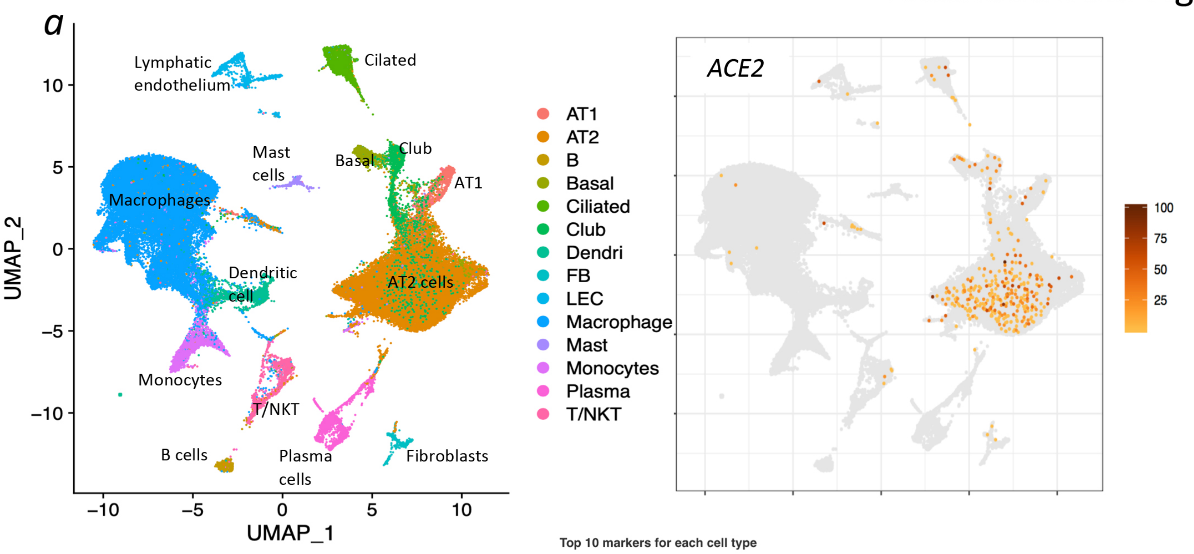


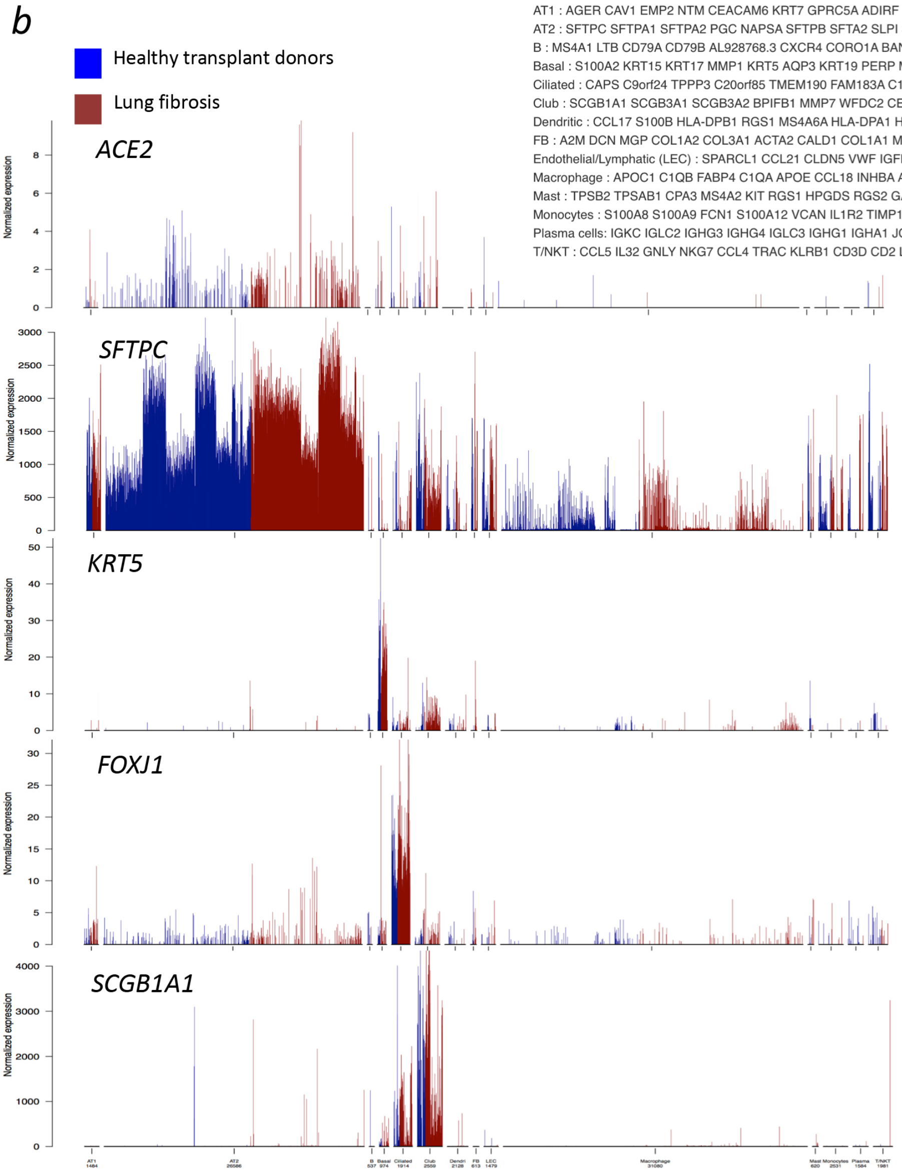


**Top 10 markers for each cell type**

AT1 : AGER CAV1 EMP2 NTM CEACAM6 KRT7 GPRC5A ADIRF SPOCK2 SCEL;
AT2 : SFTPC SFTPA1 SFTPA2 PGC NAPSA SFTPB SFTA2 SLPI SFTPD SLC34A2;
B : MS4A1 LTB CD79A CD79B AL928768.3 CXCR4 CORO1A BANK1 LIMD2 VPREB3;
Basal : S100A2 KRT15 KRT17 MMP1 KRT5 AQP3 KRT19 PERP MMP7 FXYD3;
Ciliated : CAPS C9orf24 TPPP3 C20orf85 TMEM190 FAM183A C1orf194 RSPH1 DNAAF1 TSPAN1;
Club : SCGB1A1 SCGB3A1 SCGB3A2 BPIFB1 MMP7 WFDC2 CEACAM6 PIGR MSMB LCN2;
Dendritic : CCL17 S100B HLA-DPB1 RGS1 MS4A6A HLA-DPA1 HLA-DQA1 HLA-DQB1 RNASE6 FAM26F;
FB : A2M DCN MGP COL1A2 COL3A1 ACTA2 CALD1 COL1A1 MYL9 TPM2;
Endothelial/Lymphatic (LEC) : SPARCL1 CCL21 CLDN5 VWF IGFBP7 MGP TIMP3 ACKR1 GNG11 MT1M;
Macrophage : APOC1 C1QB FABP4 C1QA APOE CCL18 INHBA ACP5 TYROBP CTSD;
Mast : TPSB2 TPSAB1 CPA3 MS4A2 KIT RGS1 HPGDS RGS2 GATA2 RGS13;
Monocytes : S100A8 S100A9 FCN1 S100A12 VCAN IL1R2 TIMP1 LST1 CORO1A CD14;
Plasma cells: IGKC IGLC2 IGHG3 IGHG4 IGLC3 IGHG1 IGHA1 JCHAIN IGHM IGHA2;
T/NKT : CCL5 IL32 GNLY NKG7 CCL4 TRAC KLRB1 CD3D CD2 LTB;

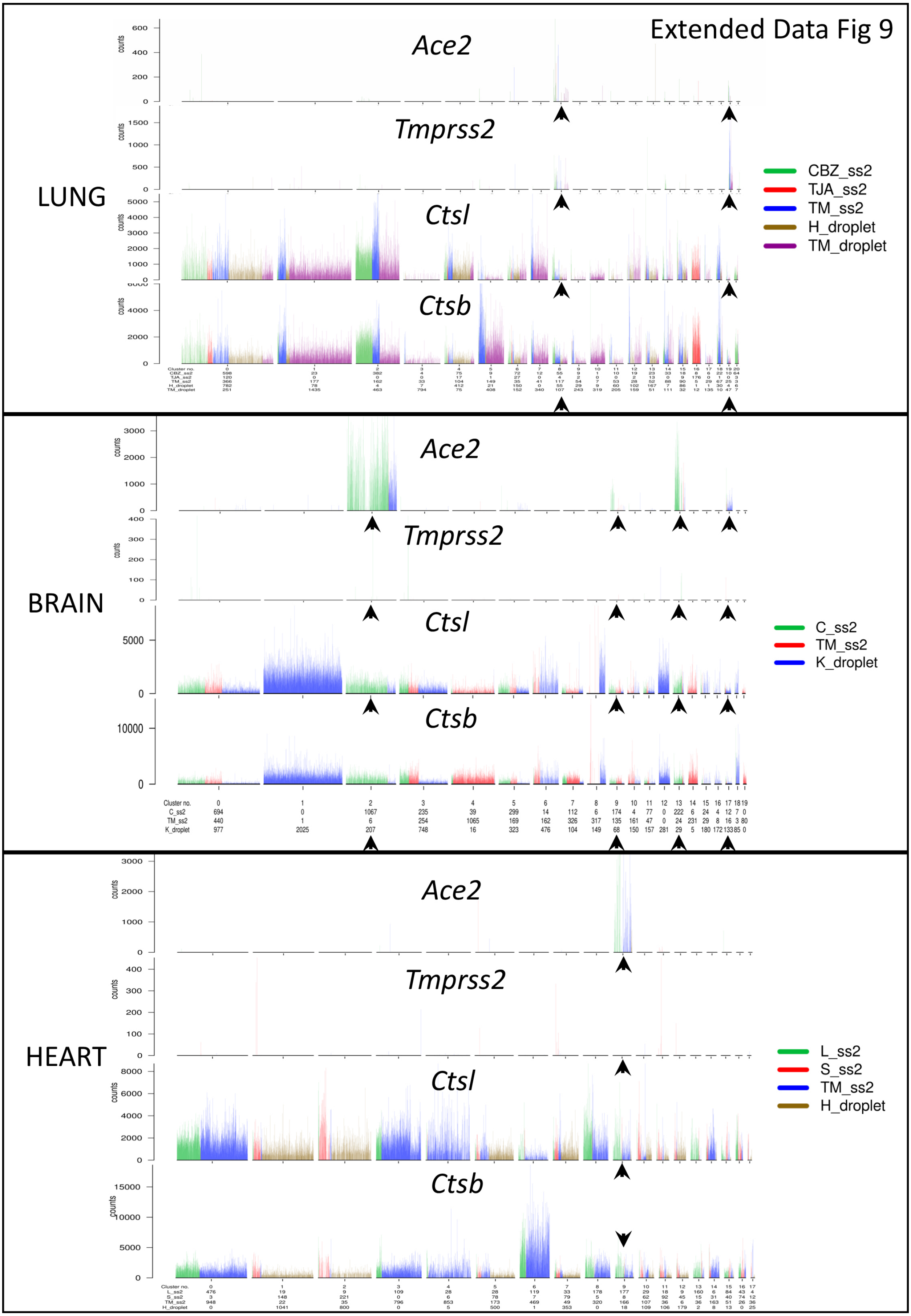
Extended Data Fig 9
LUNG
Ace2
Tmprss2
Ctsl
Ctsb
CBZ_ss2
TJA_ss2
TM_ss2
H_droplet
TM_droplet
BRAIN
Ace2
Tmprss2
Ctsl
Ctsb
C_ss2
TM_ss2
K_droplet
HEART
Ace2
Tmprss2
Ctsl
Ctsb
L_ss2
S_ss2
TM_ss2
H_droplet
counts
Cluster no.

### Extended Data Fig 10

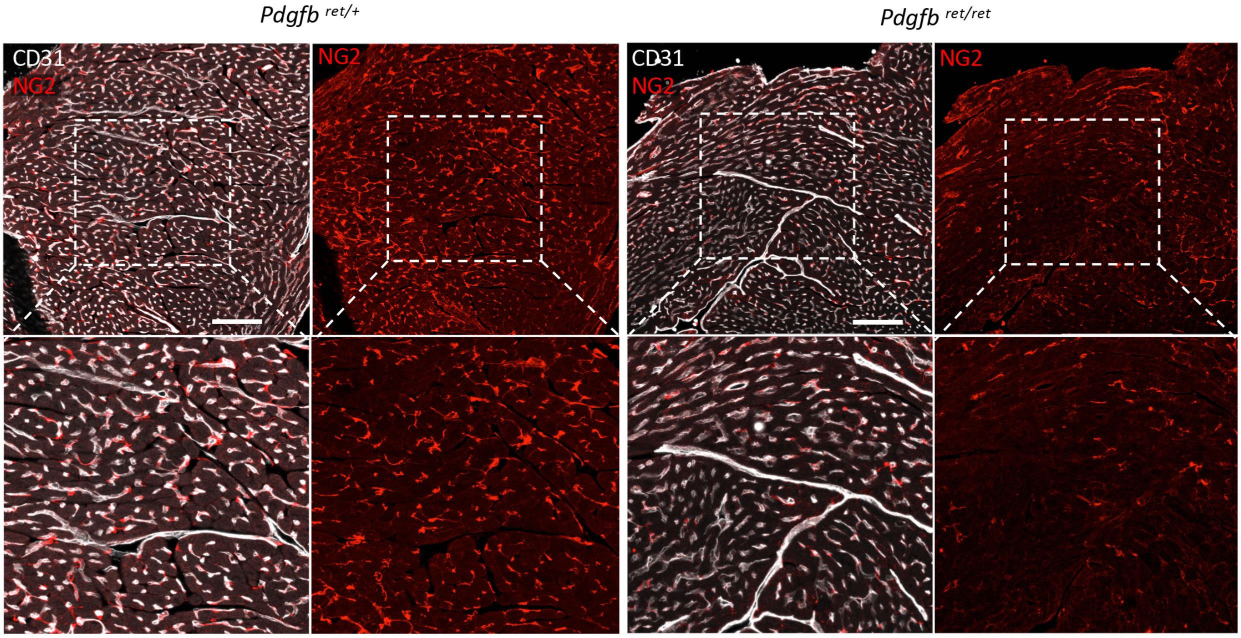

Extended Data Fig 11

*Pdgfb* ret/+ cerebral artery

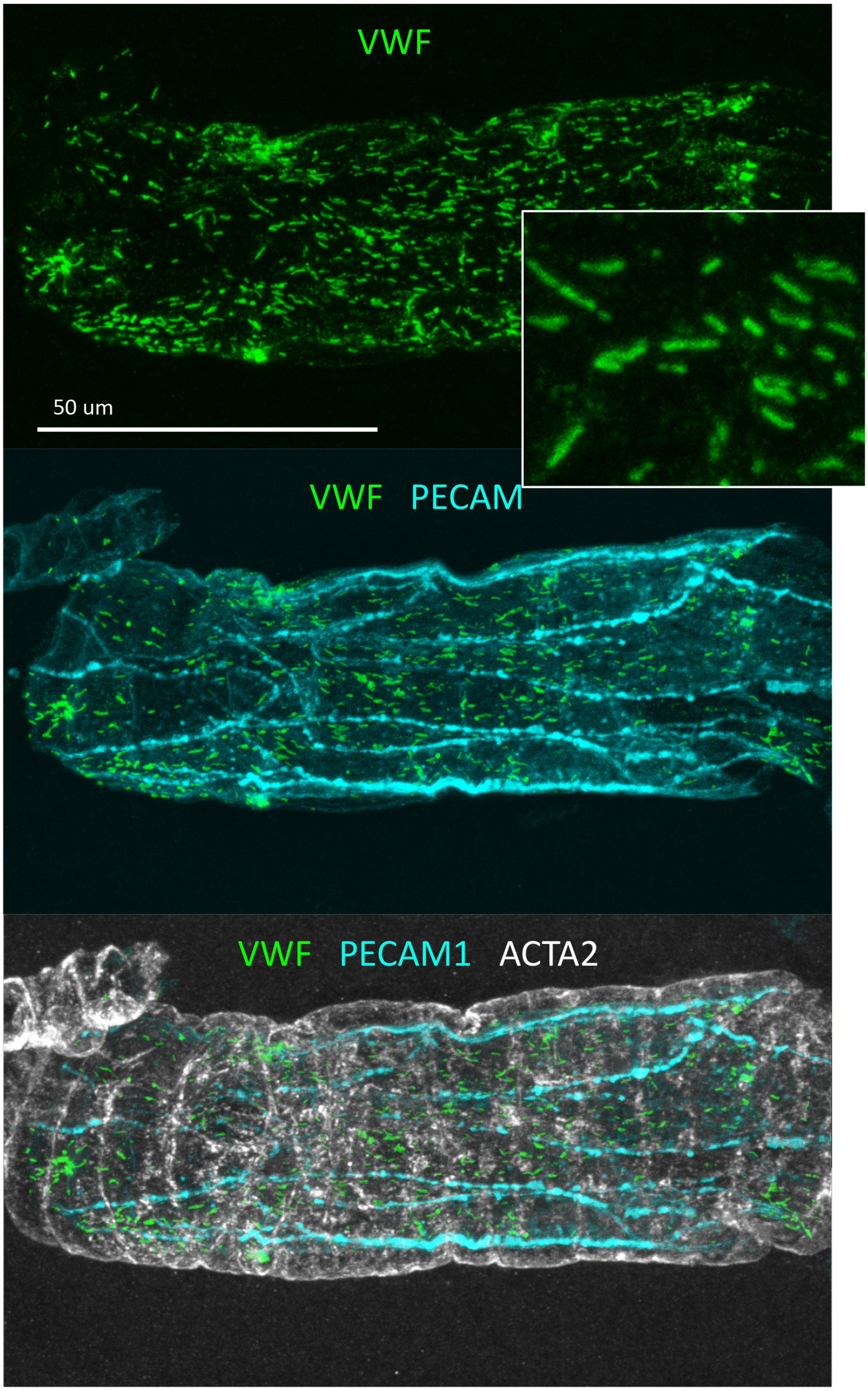

Extended Data Fig 12

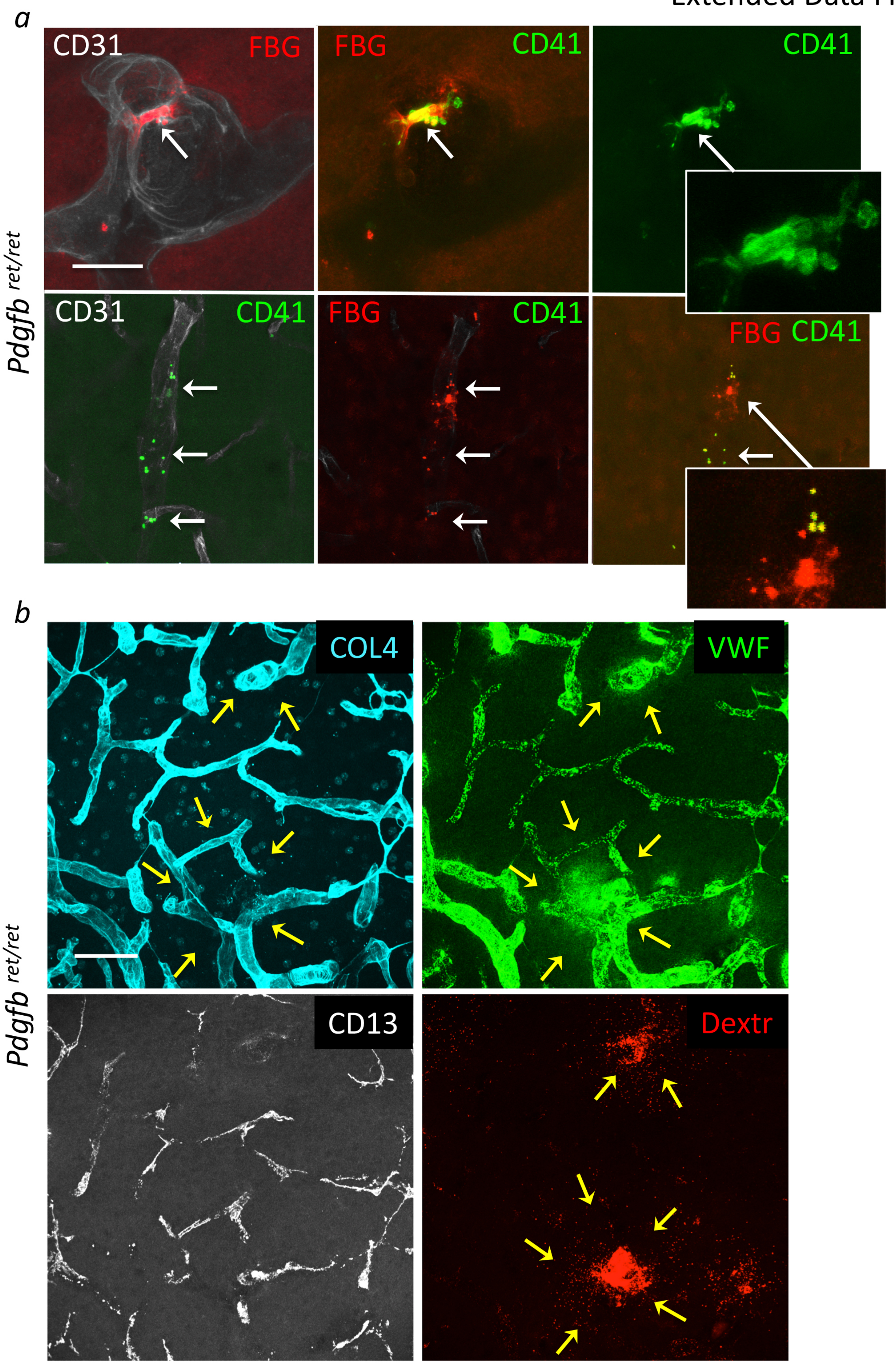

| Supplementary Table. List of used antibodies. | | | |
|---|---|---|---|
| Primary antibody | Dilutions | Supplier | Catalog number |
| PECAM1, CD31 | 1:200 | R&D Systems | AF3628 |
| PECAM1, CD31 | 1:100 | BD Pharmingen | 553370 |
| PECAM1, CD31 | 1:50 | Abcam | ab28364 |
| ANPEP | 1:100 | Bio-Rad | MCA2183EL |
| COLIV | 1:100 | Bio-Rad | 2150-1470 |
| ACTA2-Alexa Fluor 647 | 1:200 | Santa Cruz | sc-32251 |
| ACTA2-FITC | 1:200 | Sigma | F3777 |
| FIBRINOGEN | 1:200 | Dako | A0080 |
| CD41 | 1:200 | BD Biosciences | 553847 |
| PDGFRb | 1:100 | eBioscience | 14-1402 |
| VWF | 1:300 | Dako | A008229-2 |
| PODXL | 1:200 | R&D Systems | AF1556 |
| ACE2 | 1:100 | R&D Systems | AF3437 |
